# Mechanistic within-host models of the asexual *Plasmodium falciparum* infection: a review and analytical assessment

**DOI:** 10.1101/2021.03.05.434041

**Authors:** Flavia Camponovo, Tamsin E. Lee, Jonathan Russell, Lydia Burgert, Jaline Gerardin, Melissa A. Penny

## Abstract

**Background:** Malaria blood-stage infection length and intensity are important drivers of disease and transmission; however, the underlying mechanisms of parasite growth and the host’s immune response during infection remain largely unknown. Over the last 30 years, several mechanistic mathematical models of malaria parasite within-host dynamics have been published and used in malaria transmission models.

**Methods:** We identified mechanistic within-host models of parasite dynamics through a review of published literature. For a subset of these, we reproduced model code and compared descriptive statistics between the models using fitted data. Through simulation and model analysis, we compare and discuss key features of the models, including assumptions on growth, immune response components, variant switching mechanisms, and inter-individual variability.

**Results:** The assessed within-host malaria models generally replicate infection dynamics in malaria-naïve individuals. However, there are substantial differences between the model dynamics after disease onset, and models do not always reproduce late infection parasitemia data used for calibration of the within host infections. Models have attempted to capture the considerable variability in parasite dynamics between individuals by including stochastic parasite multiplication rates; variant switching dynamics leading to immune escape; variable effects of the host immune responses; or via probabilistic events. For models that capture realistic length of infections, model representations of innate immunity explain early peaks in infection density that cause clinical symptoms, and model representations of antibody immune responses control the length of infection. Models differed in their assumptions concerning variant switching dynamics, reflecting uncertainty in the underlying mechanisms of variant switching revealed by recent clinical data during early infection. Overall, given the scarce availability of the biological evidence there is limited support for complex models.

**Conclusions:** Our study suggests that much of the inter-individual variability observed in clinical malaria infections has traditionally been attributed in models to random variability, rather than mechanistic disease dynamics. Thus, we propose that newly developed models should assume simple immune dynamics that minimally capture mechanistic understandings and avoid over-parameterisation and large stochasticity which inaccurately represent unknown disease mechanisms.

## Background

The complex life cycle of *Plasmodium falciparum* (*P. falciparum*) involves many parasite stages within both the mosquito and human host. *P. falciparum* induces complex and non-sterilizing immune responses with repeated possible exposure in both the human host and mosquito. The human blood-stage infection plays a crucial role in both disease burden and transmission. Indeed, the length and magnitude of asexual parasite infection both drive clinical symptoms within a host and the transmission potential through the level of gametocytes. Thus, understanding within-host dynamics of the asexual parasite stage is essential for both the development of drugs or other tools that target asexual or gametocytes stages, and to assess burden or transmission dynamics. Within a human host, the *P. falciparum* malaria cycle begins with sporozoites transmitted by infectious mosquito bites that travel from the skin to the liver (1). Following replication in the liver, merozoites are released into the bloodstream (1,2). These merozoites subsequently infect red-blood cells (RBCs) and *in vitro,* approximately 16 merozoites emerge from a single merozoite in a 48 hour asexual blood-stage cycle (3). The blood-stage infection can persist for over 300 days if untreated (4). A small fraction of asexual parasites convert into gametocytes (2), responsible for the transmission from human to mosquito.

The asexual malaria life cycle drives clinical disease in an individual, with many processes eliciting or evading the immune response (1). After invasion of a RBC, merozoites are no longer directly exposed to immune actors. However, with hundreds of parasite proteins exported to the erythrocyte’s cell surface, immune effectors can recognize an infected RBC (iRBC) (1). Naturally acquired immune responses recognize erythrocyte surface antigens of iRBCs or antigens of the free merozoites (1), as well as antigens from liver and sexual stages (5,6). Exported parasite proteins on the cell’s surface give the cell the capability to adhere to the blood vessel’s wall, and thus evade splenic clearance (1). Furthermore, expression of the most characterized exported protein, the erythrocyte membrane protein 1 (PfEMP1), can be switched by the parasites from a large library of variants (1). New protein conformations are produced to avoid detection, requiring the host to mount new immune responses (1). *P. falciparum* escapes the immune response by successively expressing one out of 50–60 different PfEMP1 genes (7). The switching mechanisms remain uncertain, but switching between the PfEMP1 variants needs to be quick enough to evade the immune system and avoid splenic clearance, while slow enough to avoid variant exhaustion and maintain the chronic nature of the infection (7). In endemic areas where populations are continuously exposed to malaria, repeated infections lead to acquired immunity, preventing severe cases of malaria and death but without leading to sterilizing protection for infection (1).

Many mathematical and statistical models have been developed to understand population level dynamics of malaria transmission and impact of interventions (reviewed in (8,9)), or to understand within host dynamics. Although there is a long history of mathematical modeling of malaria parasite within-host dynamics over the years, the substantial biological unknown elements of both parasite and host dynamics and the highly variable nature of infection patterns, make it difficult to assess model accuracy. Furthermore, as there is limited within-host data available for infections from either immunologically naïve or non-naïve individuals, there is no “gold standard” data set or model to compare. In 1999 Molineaux and Dietz reviewed published intra-host models (10), indicating the first within-host model of malaria was likely developed in 1989 by Anderson et al. (11). Anderson and colleagues (11) described parasite dynamics via a set of differential equations representing uninfected RBCs, iRBCs, merozoites, and immune effectors (11). This model along with the others reviewed led Molineaux and Dietz to conclude that existing models lacked realism and did not make quantitative comparisons to real data. They further concluded that the reviewed models did not allow for inter-individual variability in the outputs, even though a large variation in infection dynamics exists between individuals (10). In part to address these concerns, a substantial number of mechanistic within-host models, either standalone or used in larger transmission models, have since been developed. Most of these models were initially parameterized to data from naïve patients, but not necessarily to the limited available data from previously exposed individuals.

Several sources of detailed observations of parasite dynamics and densities in naïve patients exist. In the past, malaria infection was induced to generate fever to treat other illnesses. In particular between 1917–1963 malaria was used as a therapy to treat patients with tertiary neurosyphilis before the use of penicillin (12). The most extensive malaria therapy data set was collated between 1940 and 1963 by Collins and Jeffery (12–15). The published database consists of 318 patients treated at Columbia, South Carolina and the Milledgeville, Georgia laboratories (12). This data, referred to here as the malariatherapy dataset, includes patients infected with three different strains of *P. falciparum* for neurosyphilis treatment. The data captures daily parasite counts by microscopy of both gametocyte and asexual parasites, and daily fever charts are available for each patient. This data set is the only detailed representation of the entire *P. falciparum* infection in a naïve population. Other malaria therapy data sets exist, for example of *P. vivax* infection in naïve and non-naïve individuals (16) (not further discussed here).

Over the last decade, many individual-based models of malaria transmission dynamics have been developed (reviewed in (8)); several of these include models of within-host asexual parasite dynamics (17–20). A recently published paper (21) investigated common biological assumptions made by within-host models, and concluded that current knowledge is insufficient to capture infection lengths and to explain the chronic nature of malaria infections. They further concluded their model was quite sensitive to small changes in the parameters leading to large instabilities in estimated infection lengths (21). Since asexual parasite dynamics are particularly important for modelling the effect of malaria interventions targeting humans (such as drugs or vaccines), the within-host model assumptions in these models have the potential to drive predictions at the population level, on either disease burden or intervention impact. Thus, with these within host models widely used in public health research, a good understanding of the overall dynamics, the assumptions, the uncertainties, and limitations of the models are key to critically assess and understand the predictions arising from the use of those models.

In this review, we analyzed within-host models of asexual parasitemia, and identified which components of the models drive predicted dynamics. We identified models used to investigate malaria interventions such as drugs or vaccines, either as stand-alone within-host models or used in combination with transmission models. The review and analytical assessment of each model, including the re-simulation of a subset of models to allow for deeper investigation, provides an overview of the main components of each model and their underlying assumptions. Rather than defining a gold standard, we discuss how models differ in their immune responses and parasite growth. Our comparison provides an understanding of the benefits and limitations in using these models, which directs and informs future work on within-host models of blood-stage parasitemia.

## Methods

### Models and simulation code

We identified several mechanistic within-host models currently implemented in individual-based models (IBMs) from a recently published systematic review of IBMs (8), or used as stand-alone models to understand within host dynamics or assess malaria interventions (drugs or vaccines) targeting humans. The within-host models identified as being part of malaria IBMs are Molineaux *et al.* (22), Johnston *et al.* (23), Gatton and Cheng (24), Eckhoff (19), McKenzie and Bossert (25), and Gurarie *et al.* (26). We additionally included the models of Childs and Buckee (21) and Challenger (27) (see Fig. 1, and further details on the models in *Results* and in the Additional file 1). Five of the eight identified models were re-simulated for a deeper understanding of the underlying dynamics.

**Figure 1:**
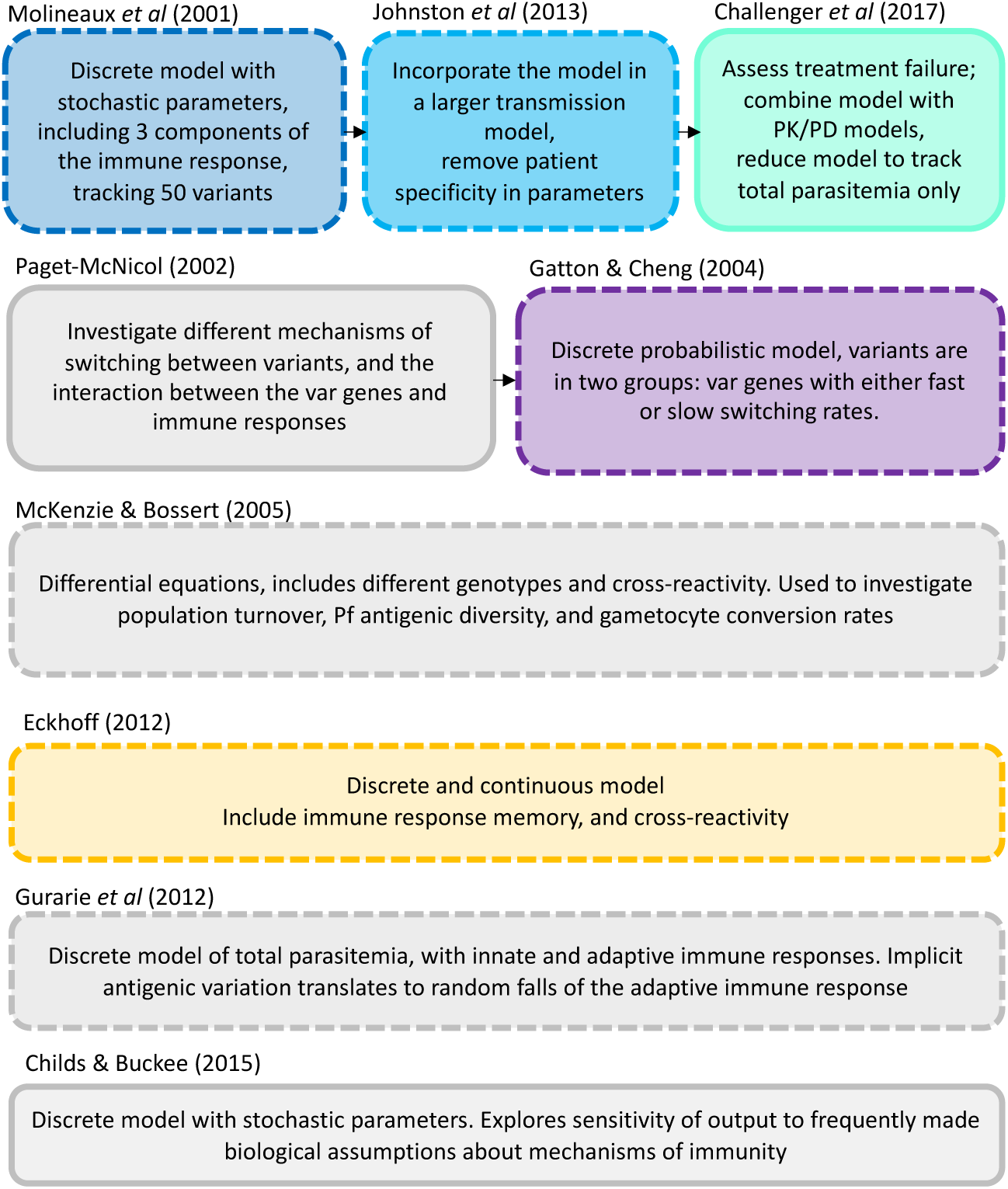
Identified within-host models of asexual parasite blood stage infection. Models are ordered by their publication date on the vertical and horizontal axis, with models adapted from another model displayed on the same row and linked with arrows. Dashed squares represent models that have been used in IBMs according to the systematic review of [6], and colored squares highlight the models that are simulated in this study, where the colors correspond to the results plotted in Fig. 3-6. Text in each box indicates a brief description of the main features or differences of the models. A more detailed description of the models can be found in the supplementary material or in the source publications (17–19,21,22,24,27)

For the simulations, we used the open access code of Johnston *et al.* (23) to recreate the Johnston *et al.* model in Matlab (28). As both Johnston *et al.* (23) and later model Challenger *et al.* (27) were based on the model of Molineaux *et al.* (22), the code of Johnston *et al.* was used as base code to reproduce the Molineaux *et al.* (22) and the Challenger *et al.* (27) models. Challenger *et al.* provides open access code for their model in C++ (29) which we used to validate our model code. The model of Gatton and Cheng. (24) was not open access but we reproduced the model in Matlab. The model code for Eckhoff (19) was available from the author’s affiliated groups (30).

### Model simulation and assessment

Each model was simulated 1750 times, which for the patient-specific model of Molineaux *et al.* corresponds to 50 simulations of the 35 malaria therapy patients used for parameterization. For all other models, this corresponds to 1750 independent realizations of the stochastic models.

Parameter values were fixed to the values defined in their respective publication (note that the parameters are either assumed from literature or fitted in their models, Additional file 1 Table S3). The same stochastic multiplication rate of each PfEMP1 variants was used in the models of Molineaux *et al.* and Johnston *et al.*, except that for Johnston *et al* any multiplication rate over 35 was resampled to be within the ranges defined by their model, as defined in their publication. Other models either had a different definition of the parasite’s multiplication rates (Challenger *et al.*) or a constant multiplication rate (Gatton and Cheng, and Eckhoff). Except for Eckhoff, our model simulations were undertaken in Matlab (28). The Eckhoff model was simulated in C++ (30). All subsequent analysis of results was performed in R (31). For the purpose of comparison, the models are all simulated for 600 days. Furthermore, we did not select for the best fit of the model to each patient and did not add measurement errors (as done by *Molineaux et al.*), as the aim was not to replicate the data but to illustrate the internal behavior of the models.

We initially assessed models via nine summary statistics for the malariatherapy data set. These summary statistics were first described in Molineaux *et al.* (22), and were later used as evaluation measures for several of the other models (23,24,27). Briefly, the nine summary statistics computed are *i)* the slope of the linear regression line from the first positive observed parasitemia to the first local maximum; *ii)* the log_10_ parasite density of the first local maximum, with a local maximum defined as a parasite density greater than the three preceding time steps (t-1 to t-3) and not lower than the three following time steps (t+1 to t+3); *iii)* the number of local maxima; *iv)* the slope of the linear regression through all the log_10_ local maxima; *v-vi)* the geometric mean and standard deviation of the geometric means between the local maxima; *vii-viii)* the proportion of positive observation in the first half and second half of the interval between the first and last positive observation; and *ix)* the last positive day (22). A positive parasite density observation was defined as an asexual parasite density equal to, or higher than, 10 iRBC per microliter. In the Gatton and Cheng model the number of iRBCs is modeled, and thus for consistency and for comparison to other models, the output was converted to iRBC per microliter assuming a body contains five liters of blood (24).

### Assessment of parasite growth and host immunity for the re-simulated models

In addition to the summary statistics, we compared estimated overall parasite multiplication rates, the innate, variant specific, general adaptive, and total immune responses, and the subsequent variation across the simulations that reflects individual variation and stochasticity. We compared models by visual inspection of time series plots of these response components and established the model’s main drivers of parasite density and infection length predictions.

To assess the modeled parasite growth, for the models which use a stochastic and variant-specific multiplication rate, we defined an average inherent parasite multiplication rate (across variants) at each time-step for each model. In Molineaux *et al.* and Johnston *et al.* each variant-specific parasite has its own multiplication rate drawn from a Normal distribution (22,23). Therefore, the average multiplication rate of all parasites at each time step is the weighted average of the variant-specific parasite multiplication rates, as follows:

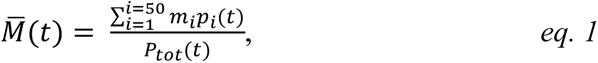

where *m_i_* is the multiplication rate of variant *i*, *p_i_*(*t*) the parasite density of variant *i* at time *t*, and *p_tot_*(*t*) the total parasite density at time *t*. For all other models, the overall parasite multiplication is an input parameter and thus does not need to be calculated.

To investigate the modeled immune responses, we categorized immune responses into innate (*S_c_*), variant-specific (*S_v_*), and general adaptive (*S_m_*). Between Molineaux *et al.*, Johnston *et al.,* and Challenger *et al.,* those three terms are directly comparable across models, but for Gatton and Cheng and Eckhoff the terms are slightly different (see Additional file 1 Table S2). Where the variant specific immune response is tracked for each variant (in Molineaux *et al.,* Johnston *et al.* and Gatton and Cheng), the overall effect of the variant-specific immune response is the weighted average of the variant-specific immune response, as follows:

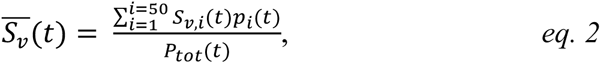

where *S_v,i_*(*t*) is the effect of the immune response on variant *i* at time *t*, *p_i_*(*t*) the parasite density of variant *i* at time *t*, and *p_tot_*(*t*) the total parasite density at time *t*. The innate and general adaptive immune responses are described as an effect on the total parasite density and are reported as such without further modifications. All equations and comparison between the models are summarized in Additional file 1 Table S2.

## Results

### Overall model structure

Via the systematic review of IBMs (8), six mechanistic within-host models were identified as part of the transmission models, as represented in Fig. 1, namely Molineaux *et al.* (22), Gatton and Cheng (24), Johnston *et al.* (23), Eckhoff (19), McKenzie and Bossert (25), and Gurarie *et al.* (26). An additional two models were identified through further literature search, namely Challenger *et al.* (27), and Childs and Buckee (21). The new quantitative results in the current study focus on five models, namely those of Molineaux *et al.* (22), Gatton and Cheng (24), Johnston *et al.* (23), Challenger *et al.* (27), and Eckhoff (19) (Fig. 1). The models of McKenzie and Bossert (25), Gurarie *et al.* (26), and Childs and Buckee (21) were included in the summary categorizations for comparison.

Most models reviewed here can generally be described in a simple discrete form

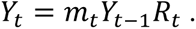

Here parasite densities or number of parasites *Y*_t_, at time *t*, depend on the parasite’s multiplication rate *m_t_* reduced by the effect of immune response *R_t_.* The immune response *R* can represent up to four components: the innate immune response, and the antibody-driven immune response defined by variant specific, cross-reactive, and adaptive immune responses, each component including different levels of stochasticity and different functions. The models can be relatively simple and reproduce a smoothed time-course of an infection, such as in McKenzie and Bossert, or include a higher level of complexity to describe more granular parasite dynamics including the typical peaks and throughs observed in clinical data. The latter usually involves explicit or implicit inclusion of a range of variant-specific parasites.

We reviewed each model via their detailed descriptions and equations in their respective publications. The models are fully described in the Additional file 1 and their main characteristics are described in Table 1, with more details of their immune response dynamics summarized in Table 2. We classified the type of equations used, the level of stochasticity integrated into the models, and detailed which dataset/s were used for calibration. We further categorized key features of parasite growth and immune dynamics for each model and summarized in tables (Table 1 and Table 2). Equations were classified and divided into four descriptions 1) parasite growth defined by the merozoite multiplication factor; 2a) triggering of, and effect of, the innate immune response; 2b) PfEMP1 variant-specific immunity dynamics; and 2c) the general or non-PfEMP1 immune response dynamic. Fig. 2 illustrates the simplified dynamics and feedback between the host and parasite for the different models. The description of the parasite growth in each model is described in Table 1 (row “Assumed multiplication rate”) and the immune responses are described in Table 2. The complete description of each model is given in Additional file 1, and a brief summary is provided here.

**Figure 2:**
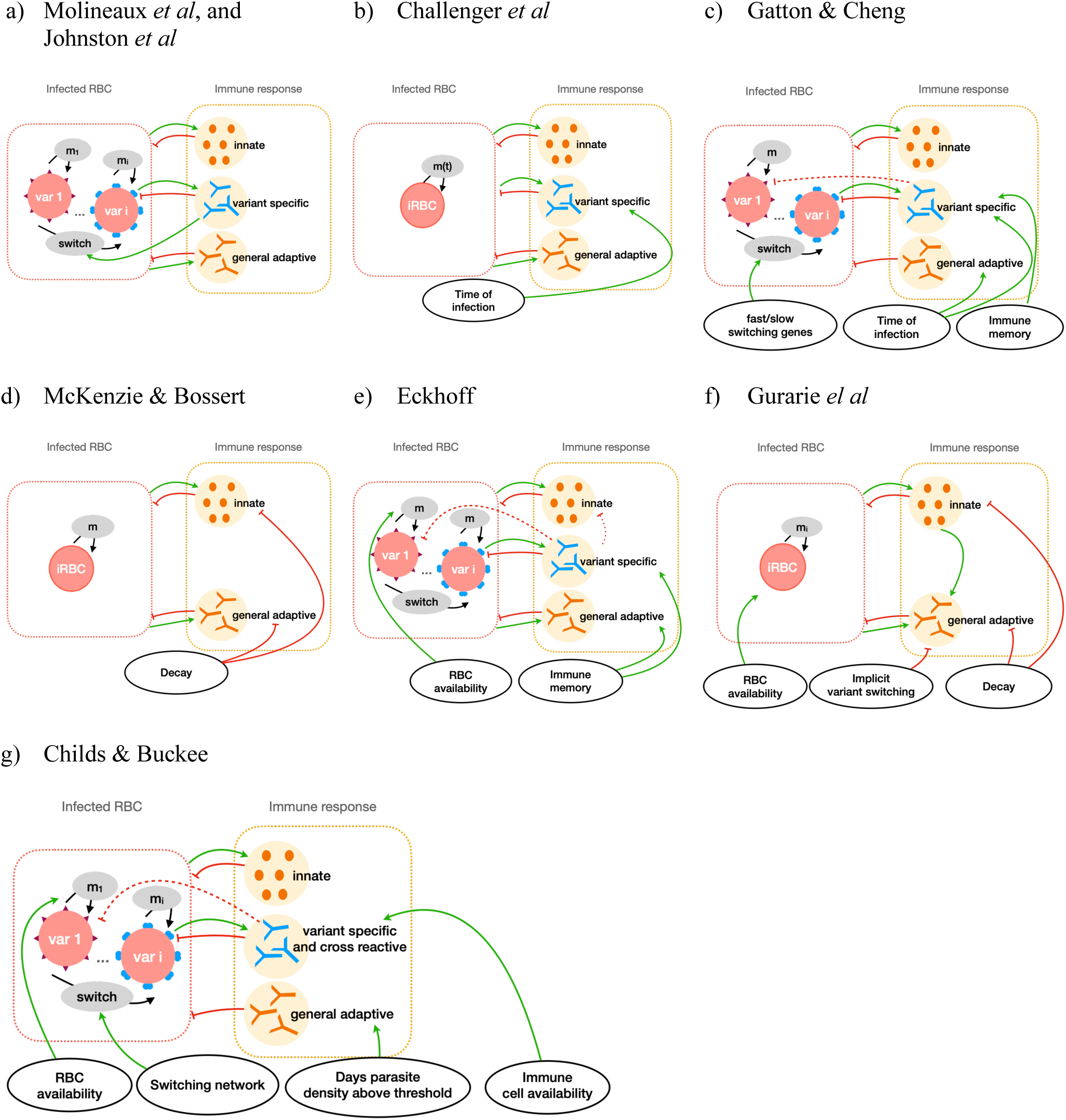
Schematic overview of the main within host dynamics. A simplified representation of the main immune and parasite dynamics for a) Molineaux et al. and Johnston et al., b) Challenger et al., c) Gatton et al, d) Mckenzie and Bossert, e) Eckhoff, f) Gurarie et al., and g) Childs and Buckee. Each model is represented by the main parasite components (left box), the host-immune component (right box) and additional factors influencing parasite dynamics (bottom circles). Models either describe overall infected red blood cells (iRBC) or total iRBC result from a sum of variant specific iRBC (var 1,..,i).Asexual multiplication (m) and variant switching of the parasite are represented by black arrows, and feedbacks between parasite and host components are represented by green arrows or red bar-headed lines for positive or negative effects, respectively. The weaker effect due to cross-reactive immune response is represented by dashed bar-headed lines. A more detailed description of the models can be found in the supplementary material, or in the source publications (17–19,21,22,24,27)

**Table 1:** Overview and main description of the models. The model description includes the main characteristics of variant or total parasite growth, the stochastic components, and the data used for fitting.

| General model structure, and parasite dynamics |  |  |  |  |  |  |  |  |
| --- | --- | --- | --- | --- | --- | --- | --- | --- |
| Models | Molineaux *et al.* | Johnston *et al.* | Challenger *et al.* | Gatton & Cheng | Eckhoff | Childs & Buckee | Gurarie *et al.* | McKenzie & Bossert |
| Adapted models |  |  |  |  |  |  |  |  |
| Publication year | 2001 | 2013 | 2017 | 2004 | 2012 | 2015 | 2012 | 2005 |
| Discrete (2 days time step) or continuous | Discrete |  |  | Discrete | Mixed | Discrete | Discrete | Continuous |
| Patient specific parameters | Yes* | No |  | No | No | No | Yes** | Yes*** |
| Tracks all parasite variants | Yes | Yes | No | Yes | Yes | Yes | No | No |
| Number of variants at the start of infection | 1 |  | - | 5 | 5 | 5 | - | - |
| Main assumption on variant switching dynamics | Dependent on variant immune response.<br>Different switching probability for each variant follows geometric distribution |  | - | Fast and slow switching variants.<br>Independent of immune response, described in (52) | Switching rate per iRBC, thus larger population have higher probability to introduce new variant. Parasites can switch to 7 available variants, out of a total of 50 variants, in each round. | Different switching networks are investigated.<br>Network assumed from (35) | Not explicit | - |
| **Assumed multiplication rates** | different for every variant but constant in time<br>($\sim \mathcal{N}(\mu_m = 16, \sigma_m = 10.4)$) | | (for overall parasites)<br>different for every time step, but correlated to previous time step | 16 (const) | 16 (const) | variant-dependent but constant in time<br>($\sim \mathcal{N}(\mu_m = 16, \sigma_m = 8)$) | range between 15-50, median 23.91 | - |
| **Additional assumptions on growth** | - | - | - | - | dependent on RBC availability | dependent on RBC availability | dependent on RBC availability | - |
| **Variability included in the model:**<br>• **stochastic parameters** | multiplication rate, patient specific parameters for the critical densities for innate and general adaptive immune response |  | critical densities for innate and general adaptive immune response, and growth rate |  |  | most parameters chosen stochastically (for sensitivity analysis) | merozoite invasion probability and replication rate, innate and adaptive immune response efficiency and activation threshold, antigenically distinct variant clusters | None |
| • **probabilistic equations** |  |  |  | effect of immune responses, antibody production, variant switching | variant switching, effect of variant specific and innate immune response |  | falls of immune response due to antigenic switch taken at random | None |
| **Data used for the fitting<sup>a</sup>** | 35 patients<sup>1</sup> |  |  | 90 patients<sup>2</sup> | Malariatherapy data<sup>3</sup> | -<sup>4</sup> | 122 malaria therapy patients | 63 malaria therapy patients<sup>5</sup> |

\* Patient specific critical densities for innate and general adaptive immune response calculation.
\*\* **Model calibrated to each of the infection chart separately to generate 50 best parameter sets for each calibrated malariatherapy data set.**
\*\*\* **Model fitted to each of the infection chart separately, thus all parameters take a patient specific value or are averaged across patients.**
<sup>1</sup> *P. falciparum* infections classified as spontaneous cures, out of a total of 334 patients in the malariatherapy data.
<sup>2</sup> Patients from the malariatherapy dataset not treated to modify the primary parasitaemia, and with no curative or subcurative treatment after the primary attack.
<sup>3</sup> Number of patients is not specified. Includes those with high initial peaks and treated during the primary peak.
<sup>4</sup> The model focuses on the effect on outcome of varying the parameters, within biologically reasonable ranges.
<sup>5</sup> inoculated with McLendon. From the patients only the days before any drug or other interventions were used, which included a range between 22 – 159 days of infection.
<sup>a</sup> as described in the initial publications, the models might have been re-fitted when updated, implemented in other research projects, or implemented in transmission models.

**Table 2:**
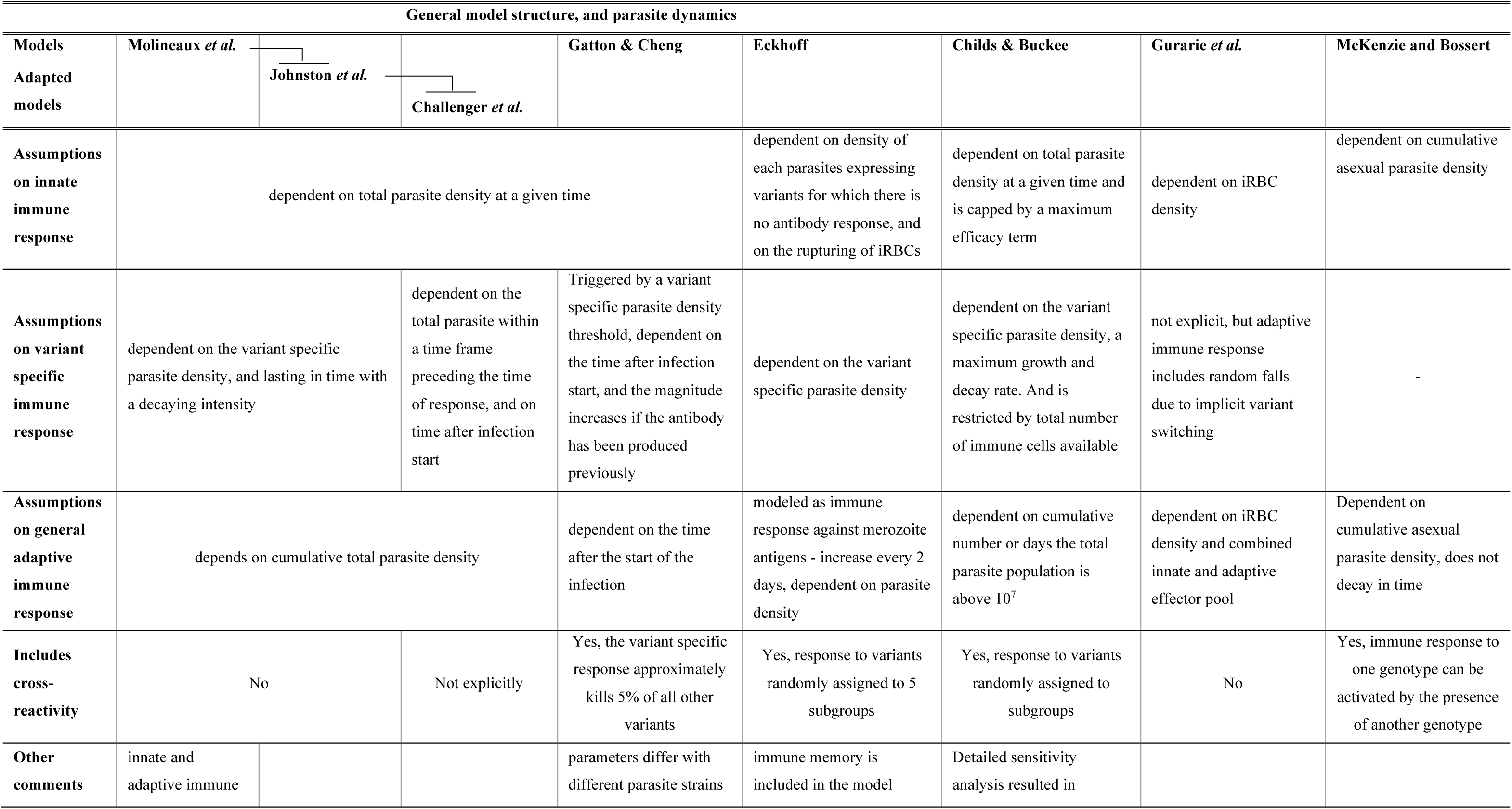
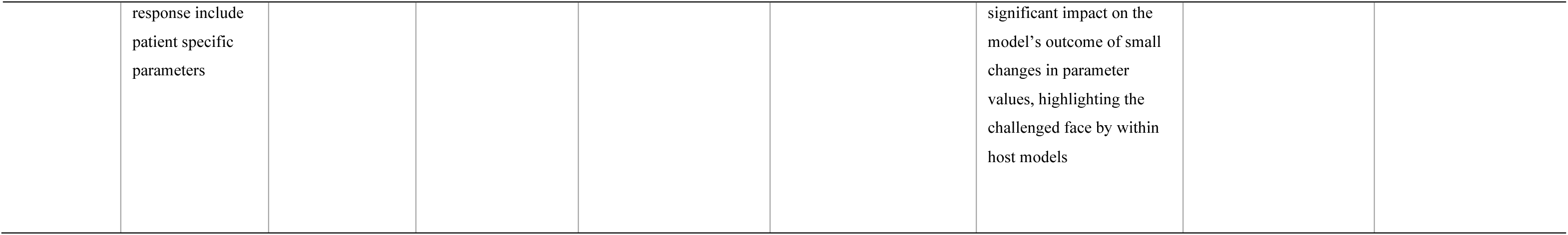
Overview and main immune dynamics of the models. The model description includes the main characteristics of innate, variant specific, and general adaptive immune response.

### Model specificities

Molineaux *et al.* describe a mechanistic malaria parasite growth model including three components of the immune response, namely an innate immune response acting early in the infection; a variant-specific immune response for each variant-specific parasite population; and a general adaptive immune response acting on total parasite population. This model was intended to reproduce parasite densities from patients in the malariatherapy dataset (12), and describes each of the 50 variant-specific parasite populations within an infection. The model of Molineaux *et al.* was adapted by Johnston *et al.* by replacing two patient-specific parameters which trigger innate and general adaptive immune response with values drawn from a distribution (see Additional file 1). Challenger *et al.* further simplified the model by tracking total parasite density instead of 50 variant-specific parasite densities to reduce memory and computational requirements. The Johnston *et al.* and Challenger *et al.* models are referred to as *Molineaux adapted* models in this manuscript.

At a similar time to the publication of Molineaux *et al.*, Paget-McNicol *et al.* (32) published a stochastic model also including three immune response components and exploring different assumptions on switching dynamics of the PfEMP1 variant expression. This model was later adapted by Gatton and Cheng (24). Instead of a set of discrete model equations with stochastic parameters describing the asexual parasite density in the blood as in Molineaux *et al*, the Gatton and Cheng model is stochastic and represent the total number of asexual parasites as a decision tree, where parasite numbers trigger choice of path in this decision tree and corresponding model equations (24). This model differs significantly from Molineaux *et al.* in regard to assumptions around the variant switching dynamics. Additionally, Gatton and Cheng include cross-reactivity between variant-specific immune responses, which was not present in the model of Molineaux *et al,* although the general adaptive immune response might capture the effect of cross-reactivity implicitly.

More recently, two additional models by Eckhoff (19) and by Childs and Buckee (21) were developed. Eckhoff’s model represents both continuous events such as immune responses to iRBC and discrete events such as the bursting of schizont and the associated immune responses. It also includes the concept of immune memory, capturing faster immune responses when an individual is re-exposed to a previously seen parasite variant. Additionally, it includes cross-reactivity between variant-specific antibody responses. In the early infection days, the probability of switching to a new variant increases with growing parasite population. The model developed by Childs and Buckee (21) is deterministic and in their work they explored a wide parameter space to assess variations in the infection dynamics with parameter choice. This model includes four different immune responses, namely innate; variant-specific; general adaptive immune responses; and similarly to the models of Eckhoff and Gatton and Cheng, includes cross-reactive immune response across parasite variants. Childs and Buckee investigate different variant switching dynamics, and explicitly limit parasite growth by available RBCs and competition of immune cells for each variant-specific immune response.

Despite these models incorporating detailed variant switching, the malariatherapy calibration datasets contain no information on gene expression profiles. Thus, there is no data describing variant expression and variant switching dynamics in the included patients. Instead, the models utilized understandings from the literature at the time of model development on potential structured switching; some of the models emphasize and discuss the uncertainty around the switching assumptions. Molineaux *et al.* and Johnston *et al* assume that immune pressure drives the switch to the expression of a new variant, based on biological studies with *P. knowlesi* infections in monkeys (7). In contrast, Paget-McNicol *et al.* refute the theory that switching depends on immunity because it can also be observed *in vitro* (33), and Gatton and Cheng’s modified model also assumes no link between variant switching mechanisms and host immune pressure. To avoid all variants being expressed with equal chances, the models either assume different switching probabilities for each variant according to a geometric distribution (as in Molineaux *et al*, and in Johnston *el al*), limit the number of available variants to switch to in each cycle (as in Eckoff), or assume two groups of variants, with either fast or slow switching behavior hypothesized at the time in (34) (as in Gatton and Cheng). The model in Childs and Buckee explores different switching networks where all variants can switch to all other variants but with different probabilities and favoring some variants over others (which was adapted from (35)).

The two last models in this review, Gurarie *et al.* and McKenzie and Bossert, are simpler models which do not include variant switching and variant-specific immune responses: thus, including only two immune response components. Gurarie *et al*. propose a discrete model that assumes the effect of the adaptive immune response is forced to have “random falls” which implicitly allow for the immune response to lose effectivity when the parasite switches to a new variant. These falls in immune response decrease in magnitude during the course of an infection, as it is assumed that the general adaptive immune response builds up. The model of McKenzie and Bossert is the only model presented here which is built around a set of differential equations in a continuous time frame. Although capable of representing the general infection pattern (high initial peak followed by a decrease in parasitemia), the model is less granular, causing model predictions to resemble a smoothed version of the malariatherapy time-series data. This model was developed to allow for different genotypes to infect the host, and to explore the effect of different gametocyte dynamics on malaria transmission. This model does not include variants nor allow for variant switching dynamics to impact immune dynamics.

### Including inter-individual variability

The models attempt to replicate time-series observed in subsets of the malariatherapy data, either with formal or less formal fitting, or at least attempting to reproduce the general pattern observed. The Molineaux *et al.* model was fitted to 35 (out of 334) patients from the malariatherapy data, who were spontaneously cured and received no other treatment. Other models used a larger number of malariatherapy patients (Table 1) to be less restrictive and avoid selection biases. More details on model fitting can be found in Additional file 1 Table S3. There is considerable variation in infection dynamics across the malariatherapy patients, both in magnitude and structure of peaks of parasite numbers, and in length of infections (Fig. 3-4). This inter-individual variability inherently includes detection and clinical measurement error but is also a result of the stochastic nature of the biological mechanisms involved. These dynamics and observed variability are challenging to replicate. In order to capture the variability, models often implement stochasticity either in the parameters defining parasite multiplication rate and/or in the implementation and effect of the innate and general adaptive immune responses (Molineaux *et al* and *Molineaux adapted*); by creating a stochastic model with most variables defined by Binomial or Normal distributions (Gatton and Cheng, and Eckhoff); or by fitting parameters separately to each patient (Molineaux *et al*, Mckenzie and Bossert, and Gurarie *et al*). Similarly, the model developed by Childs and Buckee is deterministic but can include different infection patterns by varying parameter values within the ranges specified in the publication (21).

**Figure 3:**
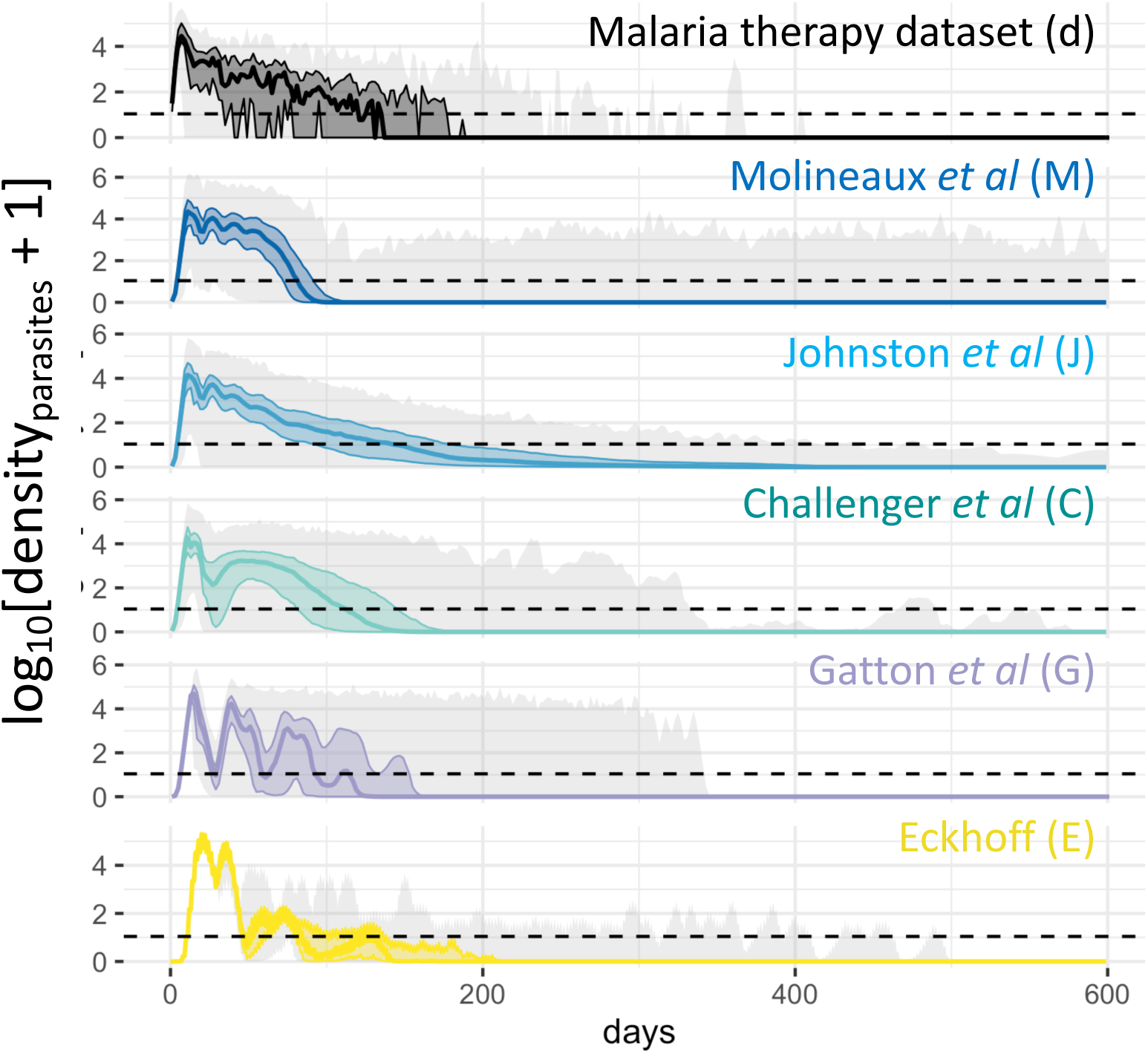
Observed and predicted total asexual parasite density. From top to bottom, the observed log_10_ asexual parasite density in time of the 35 patients from the malaria therapy dataset (d), and the predicted log_10_ asexual parasite density in time from the Molineaux et al. (M), the Johnston et al. (J), the Challenger et al. (C), the Gatton & Cheng. (G), and the Eckhoff simulations (E). The solid line represents the median across the 1750 simulations per model or 35 patients for the dataset, the colored shaded area the interquartile range (Q25-Q75) and the light grey shaded area the minimum and maximum. The horizontal dashed line indicated the threshold of positive observation (10PRBC/μl)

**Figure 4:**
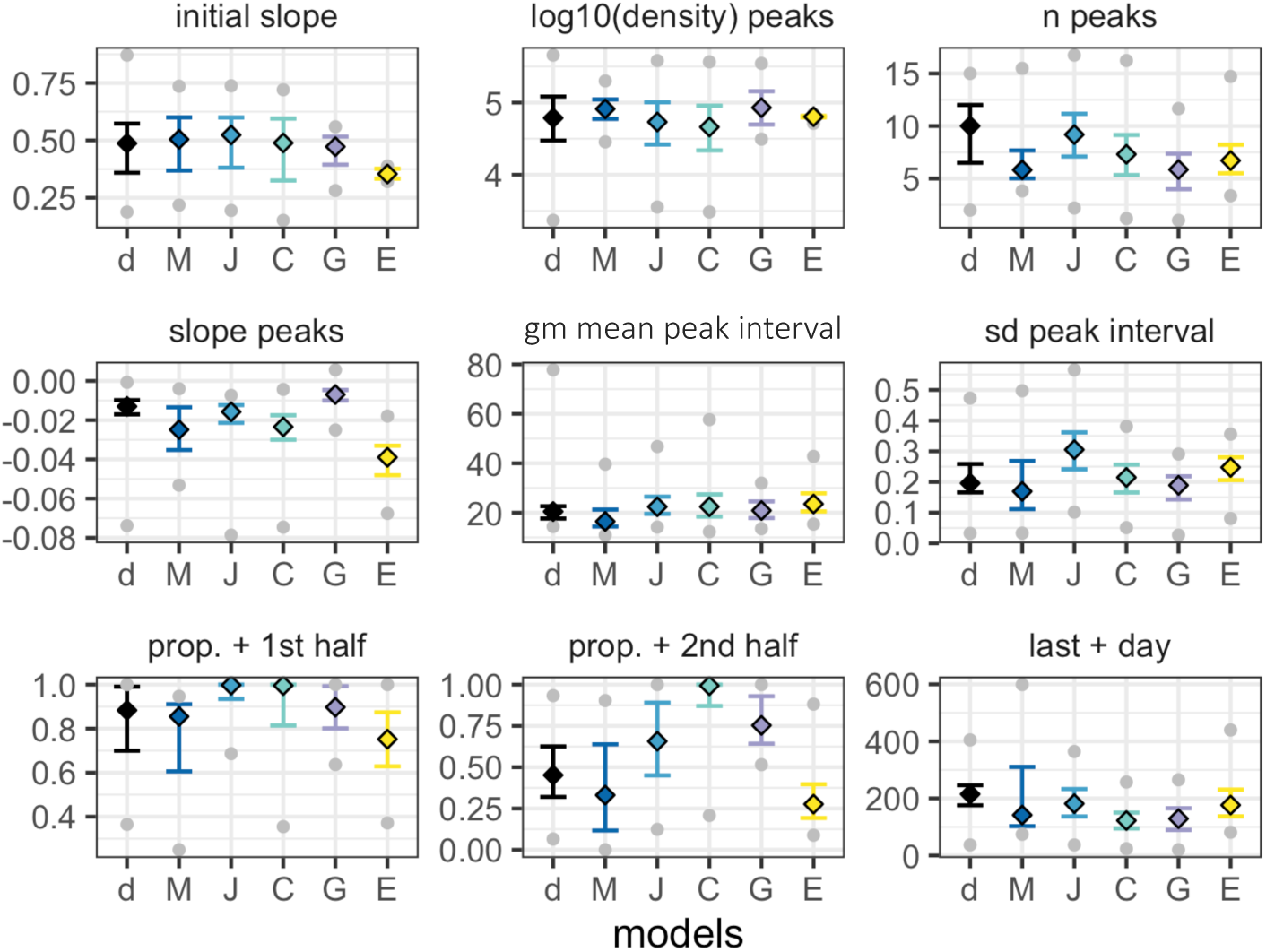
Descriptive summary statistics of the observed and predicted time course of total asexual parasite density. The nine descriptive summary statistics as defined in Molineaux et al. (see methods), for the dataset and the 5 models (by color). Squares indicate the median, bars the interquartile range and the grey dots the minimum and maximum. In each plot, from left to right, the estimates are shown for the 35 patients from the malaria therapy dataset (d), and for the simulated models of Molineaux et al. (M), Johnston et al. (J), Challenger et al. (C), Gatton & Cheng. (G), and Eckhoff (E).

### Time series of parasite density

Time series of parasite density, summary statistics (described in *methods*) of true parasitemia profiles from 35 malariatherapy patients (12), alongside time series and summary statistics calculated from 1750 simulated parasitemia profiles for the five models, are shown in Fig. 3 and 4, and in Additional file 1 Table S1. We further note that although care was taken with new or adapted code, there may be small differences from the original publications.

The observed log_10_ parasite densities of the 35 patients from the malariatherapy data varies significantly across the patients, a variability which appears to be accounted for in the models (Fig. 3). In Molineaux *et al.* most infections cease before day 200, with a subset of simulated parasitemia appearing to be chronic infections that do not end before the end of the simulation. The model of Johnston *et al.* increased the variant-specific immune response decay compared to Molineaux *et al.*, thereby decreasing the immune response’s efficiency. This results in increased simulated infection lengths compared to the simulations in Molineaux *et al.* The general shape of decay in parasite densities from Johnston *et al.* also appears to be more exponential-like compared to the other models, which is most likely a result of the more significant effect of the general adaptive immune response. In Challenger *et al.,* simulations indicate a general pattern of two peaks, the first is followed by a decrease as a result of the innate immune responses, followed by a second wave of parasitemia that is controlled by the variant-specific and general immune responses. Simulations from Gatton and Cheng indicate a slightly different time course of peaks compared to those predicted from Molineaux *et al.* and the models adapted from Molineaux *et al*. Simulations from Gatton and Cheng indicate a general first peak with a sharp decrease in parasite densities with initial innate immune responses, followed by new peaks in parasite densities given the probabilistic approach taken in Gatton and Cheng (see Additional file 1 Table S2 for the equations). The general decline in predicted parasite densities (decreasing peaks) is not reflected in the summary statistics (Fig. 4). In Eckhoff’s model the first peak is followed by much lower peaks, resulting in a steeper slope of peaks in the summary statistics (Fig. 4). The average infection lengths are consistent with the 35 malariatherapy dataset, although it is worth noting that the Eckhoff model was fitted to a larger dataset from the malariatherapy patients including more than the 35 patients in Molineaux *et al*.

### Total parasite multiplication rate during infection

We tracked each variant-specific parasite density in our simulations in all models except Challenger *et al.* and Eckhoff, the former because the equations do not explicitly model each variant and the latter due to the computational complexities of tracking each variant. The average multiplication rate at each time step is shown in Fig. 5. For Gatton and Cheng and for Eckhoff, the multiplication rate of the parasites, defined as the multiplication rate without the effect of any host immune response, is fixed at 16 per cycle and remains the same for all simulations and time (Fig.4d-e). For Johnston *et al.* and Molineaux *et al.,* each variant-specific parasite has its own multiplication rate drawn from a Normal distribution with mean of 16. For each time step, the overall multiplication rate for Johnston *et al.* and Molineaux *et al.* is thus the weighted average of the variant-specific multiplication (see *methods).* Therefore, these two models have a constant multiplication rate, in time, for each variant-specific parasite. However, the overall multiplication rate fluctuates over time, representing the competition between variant-specific parasites (with different multiplication factors), with new variants appearing, and other variants disappearing throughout the infection. For some simulations (Molineaux *et al.* and Johnston *et al.*), the overall multiplication rate is high at the beginning of the infection (Fig. 5a-b) and causes a peak in parasitemia density. The predicted infection length for a subset of simulations from both Molineaux *et al.* and Johnston *et al.* is very long, and in these subsets of simulations, the overall multiplication rate is in fact high throughout the infection (see Additional file Fig. S2).

**Figure 5:**
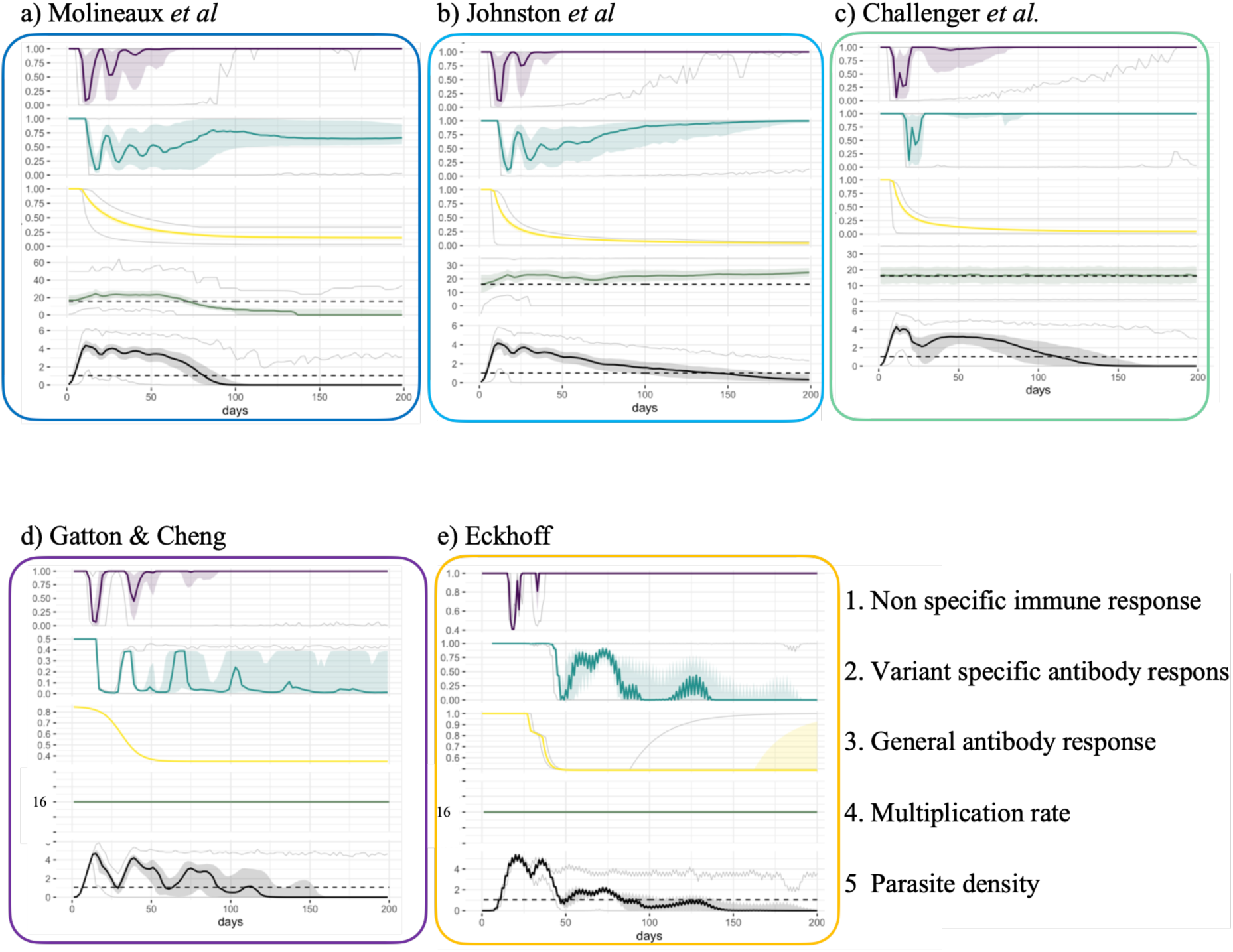
Immune responses, multiplication rates and asexual parasite densities in different models. Each panel shows, from top to bottom, 1. the effect of the innate immune response, 2. the effect of the variant specific immune response (see methods for calculations), 3. the effect of the general adaptive immune response, 4. the net effect of the product of the three immune responses, 5. the average (across variants, see methods) multiplication rate, 6. the log_10_ asexual density and 7. the product of the overall multiplication rate with the net effect of the immune response. The x-axis represents days of infection, the solid line the median, the shaded area the interquartile range (Q25-Q75) and the grey lines the minimum and maximum resulting from the 1750 simulations. The horizontal dashed green line represents an overall multiplication rate of 16, the horizontal dashed black line indicates the threshold of positive density (10PBRC/μl). Results are shown for **a)** Molineaux et al., **b)** Johnston et al., and **c)** Challenger et al., **d)** Gatton & Cheng, and **e)** Eckhoff.

In Challenger *et al.*, the multiplication rate at each time step is an input parameter of the model. It is both drawn from a Normal distribution with mean 16 and correlates with the previous time step (27) (see Additional file 1 Table S2). Thus, the median multiplication rate across all simulations remains constant at 16 by design (Fig. 5c). Taking the example of a single simulation (Fig. 6), the pattern of overall parasitemia in the Challenger model is less driven by the average growth rate in time compared with the simulations for Molineaux *et al.* and Johnston *et al.* (Fig. 6a-c). Molineaux *et al.* reported that a varying overall parasite multiplication rate, and thus the parasite inherent proliferation rate (without immunity), is essential to recreate observed peaks in parasitemia in the dataset. Furthermore, we note that long lasting infections can only occur in models that express highly multiplying variants. Overall, this indicates infection dynamics in Molineaux *et al.* and Johnston *et al.* are primarily driven by stochasticity in the inherent growth of the parasites, whereas in Challenger *et al.,* Gatton and Cheng, and Eckhoff, the stochasticity in their predicted dynamics is driven more by immune responses and subsequent killing effects.

**Figure 6:**
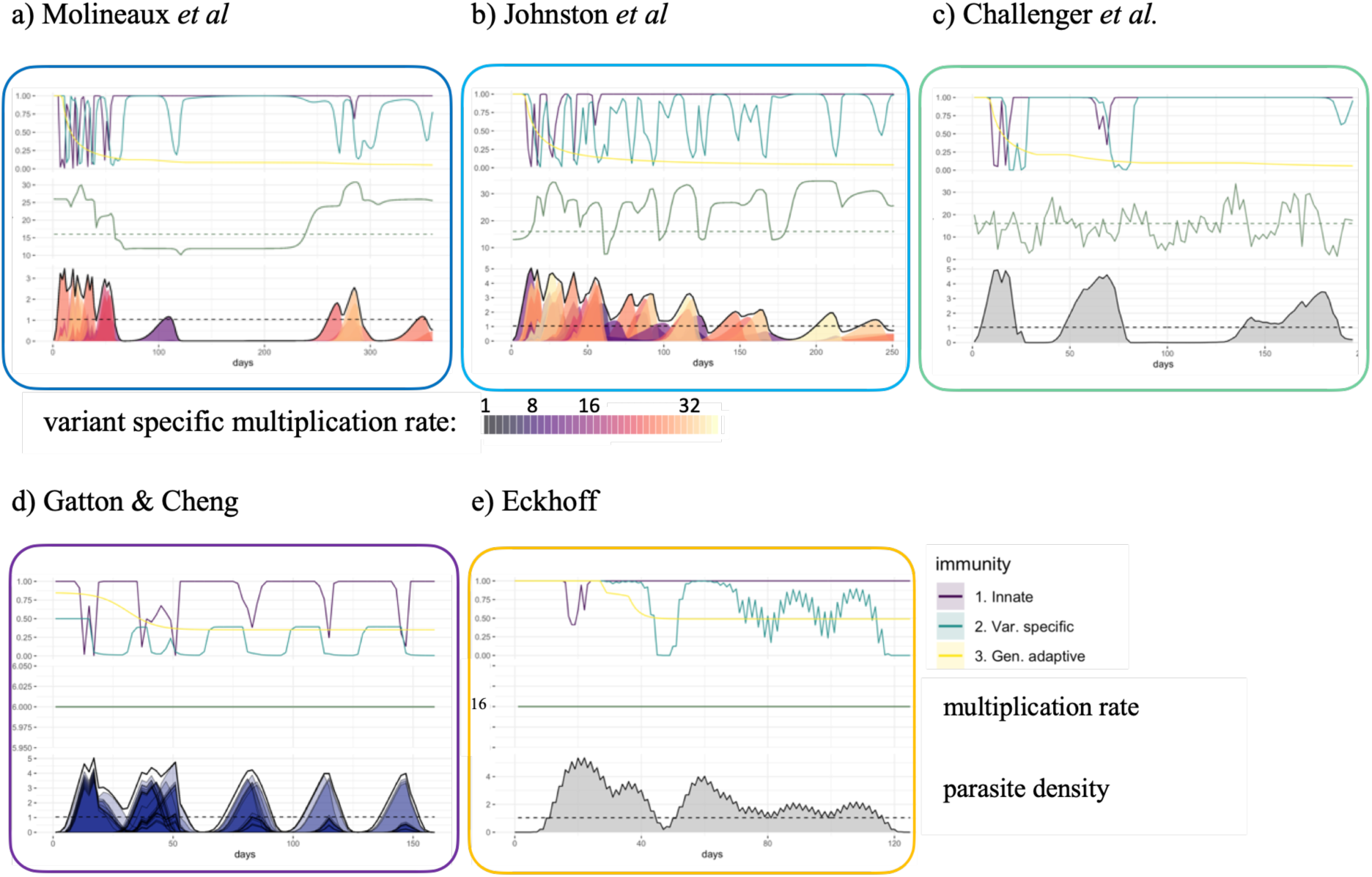
Example of the immune responses, multiplication rates and asexual parasite densities in single simulations from several models. Each panel shows, from top to bottom, 1-3. the effect of the innate, variant specific (see methods for calculations), and general adaptive immune responses; 4. the net effect of the product of the three immune responses; 5. the overall (across variants, see methods) multiplication rate; 6. the log_10_ asexual density and variant specific asexual parasite density; and 7. the product of the overall multiplication rate with the net effect of the immune response. The x-axis represents days of infection. The horizontal dashed green line represents an overall multiplication rate of 16, the horizontal dashed black line indicates the threshold of positive density (10 PBRC/μl). Results are shown for **a)** Molineaux et al., **b)**. Johnston et al., **c)** Challenger et a.l, **d)** Gatton & Cheng, and **e)** Eckhoff.

### Immune response dynamics in the models

In terms of immune response variability between the simulations (and thus assumed to exist between individuals), the highest variance across individuals arises from the variant-specific immune response, followed by the innate immune response, with the general adaptive response being reasonably consistent across individuals for the models (Fig. 5-6). The key assumptions around immune dynamics for each model are summarized in Table 2. The simulated immune responses are shown for four models in the first plots in Fig. 5, and examples of single simulations are shown in Fig.5. Note that in Gatton and Cheng and in Eckhoff, the effects of the immune responses are drawn from Binomial distributions; thus, the variability in the effects of the immune responses are not entirely captured in the immune time plots of Fig. 3 and Fig. 5.

The innate immune response is mainly active at the beginning of the infection (approximately the first 20-50 days in the different models) as it responds directly to high levels of parasite density. In Eckhoff’s model, the innate immune response is modulated by the adaptive immune response, reducing with increasing antibodies against the parasite population, making this model the only one with direct feedback between innate and adaptive immune response.

The variant specific immune response, effective against variant-specific parasite after delay, is modeled for each variant explicitly in Molineaux *et al.*, Johnston *et al.*, Gatton and Cheng, and Eckhoff. The increased decay rate in Johnston *et al.* compared to Molineaux *et al.* leads to a reduction in the immune responses (Fig. 5). In Challenger *et al.*, once the variant-specific immune response is activated, it is quickly efficient in reducing parasite densities compared to the other two models (steeper slope in the variant-specific immune response curve in Fig. 5). This efficient immune response likely explains the steep decrease after the first wave of parasite density. In Gatton and Cheng, the variant-specific response is activated when the variant-specific parasite density reaches the threshold of 12 PRBC/μl (24), and the magnitude is increased if the antibody was already produced during the infection. When there are high peaks throughout the infection, the variant-specific immune response remains very active throughout the predicted infections from Gatton and Cheng.

The general adaptive immune response increases in time since the start of the infection, responding to the cumulative parasite density (Molineaux *et al., Molineaux adapted,* Eckhoff, McKenzie and Bossert, and Gurarie *et al.* model), time (Gatton and Cheng) or a combination of both time and parasite density (Childs and Buckee). An important finding is that by comparing all three immune responses and the total net immune response (Fig. 5 and 6), the general immune response is the main effector of reducing parasite density in all models approximately after 50 days from start of infection. However, the variant immune response will ultimately end the infection. The decrease of the critical density for the general adaptive immune response in Johnston *et al.* and Challenger *et al.* compared to Molineaux *et al.* (see model description and Table S2 in Additional file 1) leads to a higher effect of the general adaptive immune response in those models, which compensates for the weaker effect of the variant-specific immune response.

## Discussion

In this study, we have reviewed eight published mechanistic within-host models of the asexual blood-stage dynamics of *P. falciparum*, of which we reproduced five. We compared predicted time-series of asexual parasitemia, modelled growth rates, innate immune responses, variant-specific immune responses, and general adaptive immune responses. The models varied widely in complexity. Whilst rather simple models such as McKenzie and Bossert have the advantage that they do not rely heavily on assumptions of unknown biological mechanisms, they reproduce less detailed patterns of infection. In contrast, more complex models such as Eckhoff or Childs and Buckee capture more detailed immune mechanisms but also include more uncertainty. Understanding the variation in multiplication rates, versus immune responses, or random effects and measurement error, and their impact on parasite density variations is particularly important when the models are included in broader investigations of the effect of a vaccine, drug or other interventions aimed to modify parasite growth patterns. The overview presented here provides a general understanding of those models.

### Model composition varies in complexity and uncertainty

Parasite and host dynamics are represented in the mechanistic models via detailed description of the parasite replication dynamics, and up to four host immune responses. Each model has its own additional complexity, specifications, and advantages. For example, to obtain increased detail of the host’s response, some models include red blood cell availability and limit the maximum immune response capacity. Or, to include more details in the parasite dynamics, some models include specific variant switching mechanisms. Models generally define a negative feedback loop between the parasite density or cumulative parasite density since the start of the infection and the effect of the immune responses, and in addition some models add the time of infection (Gatton and Cheng, Childs and Buckee, Eckhoff) as a determinant for the magnitude of the general immune response.

Including stochasticity in the models is particularly relevant for the within host dynamics of *P. falciparum* as the infection patterns observed in the malariatherapy data are highly variable among patients. Our analysis highlighted that the Molineaux *et al.* and the *Molineaux adapted* models have likely allocated too much stochasticity to the individual parasite multiplication rates, thus masking other mechanisms and placing relatively less importance on immune responses. Furthermore, for these models, we found their assumptions concerning the inherent multiplication rates of the parasites differ from other models, along with assumptions of large variability in the variant-specific parasite multiplication rates in the absence of any immune response. These dynamics were found to be essential in the Molineaux *et al.* and *Molineaux adapted* models to reproduce clinical malaria therapy patterns of infection, rather than immune responses (22). In particular, in Molineaux *et al.* and Johnston *et al.,* longer infections result from the expression of variants with a high multiplication rate towards the end of the infections for later variants. In contrast, the other models did not rely on variation to capture infection patterns. Instead, variation was mainly included in the control of infection due to immune responses and switching mechanisms.

For models including multiple parasite variants, the variant switching dynamics are an important mechanism driving the parasitemia predictions. The switching dynamics define how the parasite goes from expressing one PfEMP1 variant to another one at the next generation in order to evade the immune response. Switching dynamics in the models have been assumed to respond to the variant specific immune response (Molineaux *et al.* and Johnston *et al.*), to the current variant population size (Eckhoff), or were determined by more sophisticated switching networks (Childs and Buckee, Gatton and Cheng). In the re-simulated models, the variant-specific immune response, which is driven by the parasite’s variant switching dynamics either explicitly, or implicitly in Challenger *et al.*, was the most variable component of the immune responses across individuals. This is in contrast to the data available to inform switching dynamics over the course of an infection, which is limited to early days of infection (36–38).

The various assumptions around the mechanisms of action of the different immune responses and the interplay between parasite and host highlight the challenge of realistically reproducing the time-series observed in the malariatherapy dataset, especially the inter-individual variability. This challenge is confounded since there is limited knowledge of the biological mechanisms at play.

### Parasite multiplication rates might be lower than initially assumed

It is commonly agreed that a single iRBC produces 16 merozoites (3), of which a portion successfully invade new erythrocytes. Growth rates *in vivo* are more difficult to measure, and compared to the assumed multiplication factor of 16, includes the host-parasite interactions which reduce the observed parasite growth. Several independent statistical models previously estimated parasite growth at onset of infections in both malaria therapy or volunteer infection studies (VIS). Estimates from the malariatherapy dataset range between 10-18 (39), or 6-24 (20). In malaria VIS the growth factor was estimated to be between 12-15 (40), and in the control cohort of vaccines AMA1-based vaccine challenge between 14-21 (41). More recently, estimated *ex-vivo* multiplication factors for different malaria genotypes were found to be between 2-11 for laboratory strains and new clinical isolates (42). One hypothesis explaining the variation among parasite growth relies on the differential capacity of the PfEMP1-variants to evade splenic clearance, with the hypothesis that a subgroup of PfEMP1 expressing parasites might be fast growing (due to increased cytoadherence and thus decreased splenic clearance) (43). This mechanism may explain differing growth rates among parasites expressing different variants, and explain apparent higher multiplication rates in naïve individuals if their parasites express the fast growing PfEMP1 subgroup (43). Variance in parasite multiplication rates among clones and among infections across individuals is possible due to this variance in successful avoidance of splenic clearance during blood-stage replication, however, it is unclear whether the range should include an overall multiplication factor as large as 32 or 35, as in some models reviewed here (namely in (21–23,27)).

### Variant switching and immune response modeling are limited by current knowledge

Switching mechanisms are not well understood, and it remains unclear if the switching mechanisms are driven by antibody response (as in <u>[1], [2]</u>) or are not directly influenced by the immune pressure (as in (21,24)). In contrast to the assumptions made in the models reviewed here, it is likely that parasites express more than one variant, if not all variants, during the first blood stage generation. As highlighted by Childs and Buckee (21), this finding challenges the current understanding on the underlying mechanisms leading to chronic infections. The concept of cross-reactivity, suggested as a mechanism necessary for chronic infections (44) and included explicitly in Gatton and Cheng, Childs and Buckee, and in Eckhoff would not be applicable, as modeled and explored by Childs and Buckee, if the majority of the variants are expressed at the onset of asexual parasitemia (45). The lack of understanding about the switching dynamics supports the assumptions in Challenger *et al.*, as they only model total parasitemia without modeling switching between variants. The models described by McKenzie and Bossert, and Gurarie *et al.*, are less complex and do not model variant-specific parasitemia, offering potential modeling alternatives when detailed mechanisms of immune response are not needed. To our knowledge, there are few biological studies on the kinetics and interplay of the immune responses as defined by the models (innate, variant specific, and/or adaptive immune response). As such it is unclear how much each immune component affects the overall time course of infection. Therefore, it is not surprising that models differ in the relative importance of general or variant-specific immune responses. Moreover, in the absence of a clear biological understanding of the variability in infection time-series, model stochasticity remains an important driver of the modelled immune and parasite dynamics.

### Further data for new models

It is essential to highlight here that the malariatherapy dataset cannot be considered as a typical time course of an infection, as the data come from patients with severe neurosyphilis, who were malaria naïve, and did not include children. Furthermore, inoculations were limited a restricted set of parasite strains. Thus, although the malariatherapy data are the only detailed data available on infection time-course, models are not strictly evaluated by their ability to reproduce malariatherapy-like infections. Data on early infections are available in high detail from VIS. They provide precise quantification of parasitemia at much lower detection threshold than the malariatherapy data, and variant expression dynamics or other genetic traits relevant for understanding the dynamics in the first few days of blood stage infection can now be informed by such studies. Beyond early infections, models need to rely on longitudinal field data to assess their performance. Longitudinal field data are extremely important to explore the dynamics in realistic settings, with individuals living in endemic areas who are repeatedly exposed to malaria, including children who are most at risk for the disease, and including a range of genetic diversity and complexity of infection. Because immunity builds up with age and exposure (46), and genetic diversity is a result of immune pressure, longitudinal and cross sectional field studies which include genetic analysis give important insights in malaria infection dynamics.

The current analysis and review focused on infections in naïve individuals and we did not review the models for their ability to capture infections in pre-exposed individuals. Although data is lacking, the immune effect of pre-exposure could be added to the models as a second step, for example by adding an overall reduction factor that would lower the magnitude of the parasite density in function of age and/or exposure, similar to an empirical model by Maire *et al.* (47). The effect of co-infection was not included here and its implementation was a focus of the Childs and Buckee model (21), which hypothesizes that co-infections and super-infections have different effects based on the timing of the second infection, and that the effects of multiple infections seem to be poorly understood, and thus poorly included in models (21). Recently many field studies focusing of genetic data and analysis are giving insights in the effect and dynamics of multiple infections on a population scale (for example (48,49)), yet empirical data on the time course of complex infections are sparse and insufficient to validate models of co-infections, relying on data from mouse models for detailed infection dynamics (50,51).

## Conclusions

This review provides insights on existing models of asexual *P. falciparum* blood-stage infections, and some insights on both known and unknown biological mechanisms driving infection dynamics. Blood-stage parasite densities are at the core of malaria transmission, morbidity and mortality. Thus, population models that include models of within-host parasite dynamics to estimate the impact of blood-stage drugs or vaccine, or estimate the impact of parasite resistance, should be aware of the underlying assumptions made in the within-host model and how those changes effect infection dynamics. As in any modeling exercise, the choice of the model and its degree of complexity should depend on the specific biological, epidemiological, or public health question, and should account for available data.

## List of abbreviations

(i)RBC: (infected) red blood cell
*P. falciparum*: *Plasmodium falciparum*
PfEMP1: Plasmodium falciparum erythrocyte membrane protein 1
VIS: Volunteer infection studies

## Declarations

### Ethics approval and consent to participate

Not applicable

### Consent for publication

Not applicable

### Availability of data and materials

The outputs generated and analyzed during the current study are available from the corresponding author on reasonable request

## Authors’ contributions

Conception: FC, MAP - Design: FC, MAP - Analysis: FC, MAP, JR and JG - Interpretation: FC, MAP, TEL, LB, JR, JG - Drafted the work: FC - Substantial revision: FC, MAP, TEL, LB. All authors have reviewed and approved the manuscript

## Acknowledgements

We acknowledge and thank our colleagues in the Infectious Disease Modeling unit of the Swiss Tropical and Public Health Institute for fruitful discussions that led to improvement of this work. We thank Lauren Childs for discussions on her model in particular as well as for the other models.

## Summary of each model

### Molineaux *et al. (1)*

The model was developed to reproduce spontaneously cured malaria infection observed in 35 patients of the malariatherapy dataset who were not treated throughout the course of the therapy. The primary aim of the model was to incorporate realistic host immune responses and parasite dynamics, and to eventually include the model in a population transmission model at a later date.

The model described by Molineaux *et al.* is the first model in which immune responses against infection were divided into three components, *i)* the innate immune response, *ii)* the variant specific immune response and *iii)* the general adaptive immune response (variant transcending) (for details see Table S2). It is a discrete model with a 2-day time step to align with the merozoite’s 48 hour replication cycle, and the model allows for up to 50 variants of the parasite population. At each time step the observed parasite growth increases as a result of the intrinsic parasite multiplication rate and is reduced by the effect of the three immune responses (with each response ranging between 1 = no effect to 0 = maximum effect). Each infection begins with all 50 variants (denoted by *i* = 1,2,…,50) with initial densities set at 0, except one variant set at 0.1 PBRC/μl. At each time step parasites switch to different variants, with variant specific switching probabilities defined by a constant switching rate. The different probability of switching to each of the variants follows a geometric distribution and is modulated by the variant specific immune responses.

The innate immune response is thought to be triggered early in the infection (first few weeks), and its effect is proportional to the total parasite density at a given time, conditional on a host specific critical density. The variant specific immune response is triggered by the variant specific parasite density, effective after a certain delay (8 days), and persists with a decaying intensity. In addition, the switching mechanisms are controlled by the variant immune response, with the probability to switch to given variant increasing with high total immune responses against each of the 50 variants and decreasing with high immune response against that specific variant. The variant specific immune response, and the capacity of the parasite to switch to new variants, are thought to be a major cause for the different waves of parasitemia, and the chronic pattern of infection. The general adaptive immune response builds up slowly against the conserved part of the parasite, and acts against all variants of the parasite equally. It is effective after a delay (8 days), conditional on a host specific critical density, and responds to the cumulative parasite density with no decay in time. The general adaptive immune response is assumed to be the main cause of infection clearance, and this is confirmed by simulations except for a few simulations with chronic infection a result of stochasticity in several variant growth rates (see Fig 4 in the main).

In this discrete model, the authors include both patient specific and stochastic parameters to represent the high variability in infection dynamics among the 35 malariatherapy patients (see parameters P_m_ and P_c_, in Table S1). Variant specific parasite multiplication rates are drawn from a Normal distribution with a mean of 16 (truncated at a minimum of 1), leading to different multiplication rates for each of the 50 variants and for each simulation, but constant throughout the infection and different in each simulation. The inter-individual variability is captured by two patient specific parameters representing the critical densities for the innate immune response and the general adaptive immune response. The critical densities are calculated using the first local maximum density, and the length of the infection directly observed for each patient in the dataset.

Molineaux *et al.* additionally added a measurement error and thus were able to reproduce the 35 infections with their model by selecting, as a final step and not included in this analysis, one simulation from a pool of 50 simulations, a pool generated after trial and error, that best fit the min, median, max of the nine summary statistics, using *χ*^2^ test, for each for the 35 patients. This final step makes it difficult to reproduce the authors selection, but general infection dynamics for each patient are possible to reproduce.

### Johnston *et al. (2)*

The model was directly adapted from Molineaux *et al.* in order to be included in an individual-based model (IBM), with four parameters re-fitted via bootstrapping method to fit the minmedian and max of the nine summary statistics as well as duration o infection (more details in Table S3). This transmission model was used to understand the human component of the reproductive number R_0_, and to investigate the impact of drug treatment (through Pharmacokinetic and Pharmacodynamic (PKPD) models) and drug resistance on transmission (2). To include an adapted Molineaux *et al.* model in a transmission model, they modified the within host model to avoid the two patient specific parameters. Instead, these parameters were drawn from appropriate distributions to maintain inter-individual variability. The authors re-parametrized the patient specific parameters by using the log Normal distribution (for estimates of the first local maximum density) and the Gompertz distribution (for estimates of the length of infection) defined in previously published literature (3). To maintain the parasite multiplication rates in Johnston *et al.*within a smaller range, and thus a biologically more realistic range compared to Molineaux *et al.*, the Normal distribution was truncated not only at 1, but also on the upper end at 35.

In order to obtain a reasonable fit to the dataset following modification to the critical densities for the innate immune response and the general adaptive immune response, Johnston *et al.* re-fitted three additional parameters. All other parameters were fixed to values from Molineaux *et al*.. The refitted parameters are, *i)* the decay parameter of the acquired variant specific immune response (σ) was increased from 0.02 to 0.15, and *ii)* a constant (k_m_) allowing the calculation of the critical density threshold used for the general adaptive immune response (P_m_) was reduced from 0.04 to 0.025, and *iii)* a constant (k_c_) allowing the calculation of the critical density threshold used for the innate immune response (P_c_) was reduced from 0.2 to 0.164. By increasing the decay rate of the acquired variant specific immune response by almost ten-fold, Johnston *et al.* effectively increased infection length as an increased variant specific immunity decay rate (compared to Molineaux *et al.*) leads to a less efficient variant specific immune response. In contrast, parameters k_m_ and k_c_ in Johnston *et al.* are coupled to P_m_ and P_c_ and potentially increase the strength of the innate and general immune response (see Table S2 for full details).

### Challenger *et al. (4*)

Challenger *et al.* adapted the model from Johnston *et al.*, with the aim to assess the impact of treatment adherence and treatment failure (adding PKPD). The main adaptation of Johnston *et al.*, by Challenger *et al.* was reduction of the 50 variant specific parasite densities to a single total parasitemia, thereby reducing the number of equations from 50 to 1, simplifying the model and reducing computational time and memory requirements. The variant specific immune response in Challenger *et al.* is thus a function that represents the response to all variants instead of being the sum of responses to each variant. Early in the infection, the variant specific immune response is modeled similarly to the innate immune responses of Johnston *et al.* and Molineaux *et al.*, except for a different critical density threshold (constant for the innate immune response, stochastic for the variant specific immune response, see Table S2). During the time course of the infection, the variant specific immunity in this model responds not only to the parasite density eight days before (due to assumed delay in activation), but also to earlier time points. Thus Challenger *et al.,* in contrast to Molineaux *et al.* and Johnston *et al.*, implicitly assume immunological memory to previously expressed variants and cross-reactivity (*4*), despite not tracking all 50 variants (see Fig. S1 and Table S2).

Since Challenger *et al.* do not track 50 variants, unlike Molineaux *et al.* and Johnston *et al.*, the model considers an overall multiplication rate across all variants expressed at each time. This overall multiplication rate is computed for each time step, drawn from a Normal distribution around 16 while being positively correlated to the previous time step, implying that this stochastic, but correlated, parasite multiplication rate is the result of the emergence and disappearance of variants during the time course of an infection (4).

The model of Challenger *et al.* was fitted to the same 35 malariatherapy patients as in Molineaux *et al.,* fitting the parameters of the overall variant specific immune response and the overall multiplication rate, and re-fitting the constant allowing the calculation of the critical density threshold for the general adaptive immunity (k_m_). The authors used Markov chain Monte Carlo method u(Metropolis-Hasting algorithm) to generate random walks in parameter space and fitted the model to the nine summary statistics (see Table S3). A constant that allowed the calculation of the critical density threshold for the general adaptive immunity (k_m_) was reduced from 0.0250 to 0.021 which leads to a slightly stronger general adaptive immune response in Challenger *et al.* compared to Molineaux *et al..* Additionally, the critical density threshold of the innate immune response function (P_c_) was drawn from a log Normal distribution with σ = 1.2 instead of σ = 1.148 as in the other models. All other parameters are identical to the model in Johnston *et al.*.

### Gatton & Cheng (5)

Gatton & Cheng (5) is a model adapted from a previously published probabilistic model of Paget-McNicol *et al.* (6). The model was fitted to the malariatherapy data differentiated by the *P. falciparum* I strain used. The model was fitted using sets of 100 simulations, altering 6 parameters in the model. They compared the model output to four variables, namely maximum parasitemia, number of days with fever, number of days with parasitemia higher than 10/microliter, and number of days with parasitemia higher than 10000/microliter (see Table S3).

As with Molineaux *et al.*, it tracks 50 different variant specific parasites, starts the infection with only one variant expressed, tracks the parasites in 2-day time steps, and includes three different types of immune responses. The model describes the total number of asexual parasites via a flowchart which acts more like a decision tree with triggers different equations for given parasite numbers (5) (see equations in the Table S2). At each time step the parasites undergo replication with a multiplication rate of 16, and the probability of success of the replication is computed from Binomial distribution dependent on the general adaptive and variant specific immune response. Thus, in the absence of any immune response, the parasites have a constant growth rate of 16.

The innate immune response in Gatton & Cheng is activated after reaching a threshold and is assumed to increase with a higher number of parasites until a maximum effect is reached. The effect of the variant specific immune response is modeled as being inversely proportional to the number of antibodies produced, the latter being computed at each time step. The magnitude of the antibody response is dependent on whether this antibody has previously been produced during the infection, as well as cross-reactivity (so that an antibody response against a specific variant also has a small effect on all other variants). The general adapted immune response is only dependent on the time of infection, thus not changing with parasite numbers, nor does it differ between individuals (see Table S2). Additional biological differences between the model of Gatton *et al.* compared to Molineaux *et al.* and Johnston *et al.* is the switching mechanism from one variant to another, which in Molineaux *et al.* and Johnston *et al.* is assumed to depend on the variant specific immune response, whereas in Gatton & Cheng it is completely independent from the host’s immune response. In addition, Gatton & Cheng assumes two populations of variants, either fast or slow switching, with 10 variants considered fast switching (rate ∼ 0.5-4.5%) and 40 in the slow switching group (rate ∼ 9.10-11.8 %), compared to 0.02% in Molineaux *et al.* and Johnston *et al.*. Switching dynamics were previously investigated by the author (7) and based on *ex-vivo* analysis of var gene transcripts of parasites isolated two volunteers who underwent challenge infection (8).

### Eckhoff (9)

A summary and illustration of this model can be found on (10). This model has both discrete and continuous components, to represent discrete events such as schizont rupture in a two-day interval and the antibody response against merozoites, and to represent the continuous immune components acting on infected red blood cells (iRBCs). This model is more complex in the immune response mechanisms described, compared to the Molineaux *et al.* and Molineaux adapted models, as it explicitly includes antibody responses to PfEMP1 and with smaller effect to minor epitopes, it includes immunological memory of the antibody response, and it includes continuous and discrete immune responses.

The blood-stage immune response is divided into four components, namely the innate immune response, the antibody immune response against each PfEMP-1 variant, and each shared minor epitope, and the antibody immune response against merozoite proteins (such as MSP-1 and AMA-1). As for the Molineaux *et al.* and Molineaux adapted models, the innate immune response is assumed to limit the maximum parasite density (first peak). This innate immune response acts against both iRBCs (continuous component) and against ruptured schizonts (discrete component). Its effect is activated by a pyrogenic threshold, and it decreases with increased antibody levels. The antibody response is dependent on the capacity to generate specific antibodies, which depends on the concentration of the variant specific parasite and on the immune memory to the parasite. Similarly, to other models, the antibody response decays once the corresponding parasite population is absent, but the antibody production capacity keeps an immunological memory. Thus, once exposed again to the same variant, the antibody response will reach higher levels faster. The immune response against merozoite proteins has a similar effect as the general adaptive immune response in the other models. It increases in time and leads to a decrease of the parasite peaks amplitude in time. The PfEMP1 variant specific immune response eventually clears the infection.

Initially, the model was fitted to malariatherapy data, but contrary to Molineaux *et al.,* the dataset includes patients with reinfection (as memory of immune response is modeled) and the perturbed ones to avoid selection biases. Parameters are either assumed through literature, or fitted to the malariatherapy dataset (details in (9)). The model outputs were compared to the first parasite density peak, lower secondary peaks, and lower peaks after 100 days, the possibility of reinfection with homologous strains, the interval between peaks, and the distribution of measured durations (9). The model has been re-fitted more recently using field data (11), and re-calibrated parameters included the number of PfEMP1 variants the switching rate, the number of MSP variants in the overall parasite population, the fraction of merozoites inhibited by maximum MSP1-specific antibody level, the number of minor epitope variants, and the kill rate of infected red blood cells due to antibody response to minor epitopes. As described in (11), a Dirichlet-multinomial distribution was used to compare simulation data with field data (12,13).

### Childs & Buckee (14)

This is a deterministic discrete model with 2-day time steps. It tracks parasite population of 60 variants and includes four different immune mechanisms. The immune response includes innate, variant specific, general adaptive immune response, similar to the other models, as well as a cross-reactive immune response. Parasite growth depends on the variant specific inherent multiplication rate, drawn from a Normal distribution, and parasite growth is limited by red blood cell availability. Similar to the other models, the innate immune response depends on the current number of parasites, the variant and cross-reactive immune response depend on the variant specific (or group of variants for the cross-reactive immune response) parasites, and the general immune response grows slowly during the course of an infection. Differing to the other models, the four immune responses are all capped by a maximum efficacy level, and the variant specific and cross-reactive immune response depend on the number of available immune cells. The cross-reactive immune response acts on groups of variants randomly grouped and pre-determined. The general immune response, instead of being dependent on time or cumulative parasite density like the other models, here increases each day that the number of parasites are above a certain threshold.

Because the main aim of this model was to understand the key assumptions of the biological mechanisms included in the mechanistic within host model, the model’s main advantage compared to the others rely on the sensitivity analysis in the parameter values. Indeed, it is a deterministic model, but a wide range of values for all the parameters have been evaluated using the Latin Hypercube Sampling method. In addition, different switching networks, enabling parasites to switch from one variant to another, were evaluated.

### Gurarie *et al.* (15)

This is a discrete 2-day time step model. In this model there is a continuous production and loss of uninfected RBCs, which can become infected by the release of merozoites, and parasite replication is inhibited by two immune effectors, innate and adaptive. The merozoite invasion depends on available RBCs so that when the ratio between the number of merozoite and the number of available RBCs becomes large, merozoites start competing for available RBCs. Despite this assumption, it is not clear RBC availability is important for human *P. falciparum* infection. The innate immune response depends on the number of infected cells, activated after reaching a certain parasite density threshold level, and reaching a maximum clearance level. The adaptive immune response is triggered by the product of infected RBCs, and the combined effector pool of the innate and adaptive response, which is a way to allow for adaptive immune memory, and it also depends on an activation threshold. Similar to the other models, the adaptive immune response takes more time to develop but has a slower decay time compared to the innate immune response.

Rather than explicitly modelling the different variant populations, the variant switching dynamics, and variant specific immune response, such as in other models, the model greatly simplifies the equations by using proxies of the more complex underlying mechanisms. First the model only includes innate and adaptive immune responses, making no distinction between the variant specific and general adaptive immune response. Second the parasite population is taken as a whole and does not differentiate between the variant-specific populations. Nevertheless, the variant switching and resulting decrease in an effective adaptive immune response is implicitly accounted for by forcing “random falls” at each replication cycle (where potentially parasites switched to a new variant) of the effective adaptive immune response. To implicitly account for the variant-transcending immune response, the amplitude of the random falls of the effective adaptive immune response decrease in time.

The model was fitted using the malariatherapy data (from 122 patients), and each malariatherapy patient was considered a random realization of a stochastic process. The fitting process was done in two steps to fit the first parasitemia peak assumed deterministic and a second stochastic step for parasitemia post first peak. The model outcome was compared to minimum and maximum of parasite density time-series for each patient, and 50 simulations per patient were selected as an ensemble of best choices (Table S3). The calibration process of the model resulted in a range of parameter values that can be randomly chosen at each realization, and thus allow for variability in the parasite dynamics as observed in the malariatherapy dataset. These parameters relate to the invasion probability, the replication of parasites, immune response efficiencies and activation thresholds, and the assumed antigenically distinct variant clusters.

### McKenzie & Bossert (16)

This model is the only model studied here that uses differential equations rather than discrete equations to represent the malaria within host dynamics. The model includes merozoite replication and conversion of parasites into gametocytes and includes the effect of two types of immune responses, innate and adaptive. There is no distinction of variant specific parasites. Both immune responses are stimulated by the asexual density, with a stronger stimulus of the innate effectors, but the dynamic of the innate effectors assume the decay and the removal of the effectors, while the adaptive immune response grows with the asexual parasite density and has no decay or removal in time (note that the equation for the adaptive effectors include a decay component, but its parameter *w* = 0 in (16).).

Although not explored in this study, the model tracks different genotypes, includes cross-reactivity, and in addition, investigated several aspect of gametocyte dynamics. Cross-reactivity is included by allowing for immune responses to one genotype to be stimulated by the asexual parasite density of another genotype.

The model was fitted to data from 63 malariatherapy patients who were inoculated with the McLendon strain, including only the parasite densities in time before any drug was administered. The model was fitted to data from each of the 63 patients individually, using the steepest-descent algorithm and comparing model outputs to parasite density time-series (Table S3), and an average of the parameter values was reported and used for their transmission model.

---

**Figure S1:**
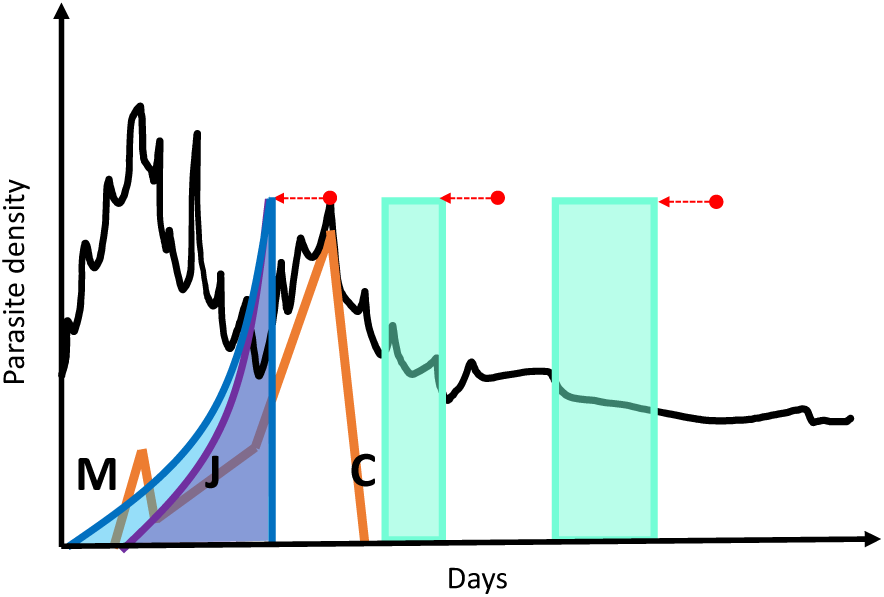
Schematic illustration of an infection and variant specific immune responses at a given time point from models Molineaux et al., Johnston et al., and Challenger et al.. Example illustration of total parasite density in time (black) with an example of a single variant parasite density in orange. The parasite density affecting the variant specific immune response is represented by the coloured area in blue for Molineaux et al., in purple for Johnston et al., and in green for two examples at two different time points for Challenger et al.. At a given time point, represented by a red dot, the variant specific immune response in the Molineaux et al. (M) and Johnston et al. (J) model depend on the variant specific parasite density (orange) from eight days prior to the time point, and the effect of the parasite density decreases in time, with Molineaux et al. assuming a slower decay rate than Johnston et al.. For Challenger et al. (C), the total parasite density (black) in a given time frame eight days prior to the time point affect the variant specific immune response, with the time frame increasing further along in the infection, implicitly including immune memory, here represented by an example of two time points (red).

**Table S1:** Summary statistics of the observed malariatherapy data and the models. Summary statistics from the simulation are shown in the first row, and the published results in the second row in italics. Source of published statistics: data [1], Johnston et al.(2), Challenger et al.(4), Gatton & Cheng (5).

|  | Data |  |  | Molineaux *et al* |  |  | Johnston *et al* |  |  | Challenger *et al* |  |  | Gatton & Cheng |  |  | Eckhoff |  |  |
| --- | --- | --- | --- | --- | --- | --- | --- | --- | --- | --- | --- | --- | --- | --- | --- | --- | --- | --- |
|  | med | min | max | med | min | max | med | min | max | med | min | max | med | min | max | med | min | max |
| initial slope | **0.49** | 0.19 | 0.87 | **0.50** | 0.22 | 0.74 | **0.52** | 0.19 | 0.74 | **0.49** | 0.15 | 0.72 | **0.47** | 0.28 | 0.56 | **0.35** | 0.32 | 0.39 |
|  |  |  |  | *0.49* | *0.19* | *0.87* | *0.52* | *0.24* | *0.75* | *0.49* | *0.25* | *0.71* |  |  |  |  |  |  |
| log10(density) peaks | **4.79** | 3.37 | 5.66 | **4.91** | 4.46 | 5.30 | **4.73** | 3.55 | 5.58 | **4.66** | 3.49 | 5.56 | **4.93** | 4.49 | 5.54 | **4.80** | 4.76 | 4.84 |
|  |  |  |  | *4.79* | *3.37* | *5.66* | *4.78* | *3.69* | *5.67* | *4.77* | *3.57* | *5.61* | *4.81* | *4.47* | *5.31* |  |  |  |
| n peaks | **10.00** | 2.00 | 15.00 | **5.83** | 3.83 | 15.49 | **9.19** | 2.20 | 16.74 | **7.31** | 1.20 | 16.23 | **5.86** | 1.03 | 11.66 | **6.71** | 3.37 | 14.54 |
|  |  |  |  | *10.00* | *2.00* | *17.00* | *9.00* | *2.00* | *17.00* | *7.70* | *1.70* | *15.50* | *10.00* | *1.00* | *27.00* |  |  |  |
| slope peaks | **-0.01** | -0.07 | 0.00 | **-0.02** | -0.05 | 0.00 | **-0.02** | -0.08 | -0.01 | **-0.02** | -0.07 | 0.00 | **-0.01** | -0.03 | 0.01 | **-0.04** | -0.07 | -0.02 |
|  |  |  |  | *-0.01* | *-0.07* | *0.00* | *-0.02* | *-0.09* | *-0.01* | *-0.01* | *-0.03* | *0.00* | *-0.01* | *-0.03* | *-0.01* |  |  |  |
| geo. mean peak interval | **20.48** | 14.40 | 77.77 | **16.47** | 10.95 | 39.61 | **22.40** | 14.19 | 46.77 | **22.41** | 12.31 | 57.67 | **20.89** | 13.55 | 31.97 | **23.56** | 15.51 | 42.45 |
|  |  |  |  | *20.00* | *14.40* | *77.80* | *14.60* | *1.50* | *28.40* | *22.80* | *12.20* | *61.00* | *22.00* | *14.20* | *29.30* |  |  |  |
| sd peak interval | **0.20** | 0.03 | 0.47 | **0.17** | 0.03 | 0.50 | **0.31** | 0.10 | 0.57 | **0.21** | 0.05 | 0.38 | **0.19** | 0.03 | 0.29 | **0.25** | 0.08 | 0.35 |
|  |  |  |  | *0.20* | *0.03* | *0.47* | *0.31* | *0.04* | *0.56* | *0.22* | *0.03* | *0.48* | *0.22* | *0.07* | *0.29* |  |  |  |
| prop. + 1st half | **0.88** | 0.37 | 1.00 | **0.86** | 0.25 | 0.95 | **1.00** | 0.69 | 1.00 | **1.00** | 0.35 | 1.00 | **0.90** | 0.64 | 1.00 | **0.75** | 0.37 | 1 |
|  |  |  |  | *0.88* | *0.40* | *1.00* | *0.97* | *0.57* | *1.00* | *0.97* | *0.46* | *1.00* | *0.67* | *0.48* | *1.00* |  |  |  |
| prop. + 2nd half | **0.45** | 0.07 | 0.93 | **0.33** | 0.00 | 0.90 | **0.66** | 0.12 | 1.00 | **0.99** | 0.21 | 1.00 | **0.75** | 0.52 | 1.00 | **0.28** | 0.09 | 0.88 |
|  |  |  |  | *0.46* | *0.08* | *0.94* | *0.58* | *0.12* | *1.00* | *0.79* | *0.10* | *1.00* | *0.61* | *0.37* | *1.00* |  |  |  |
| last + day | **215** | 37 | 405 | **142** | 74 | 599 | **182** | 37 | 366 | **123** | 24 | 257 | **129** | 20 | 265 | **177** | 82 | 439 |
|  |  |  |  | *215* | *37* | *405* | *193* | *38* | *404* | *198* | *39* | *409* | *239* | *10* | *590* |  |  |  |

**Figure S2:**
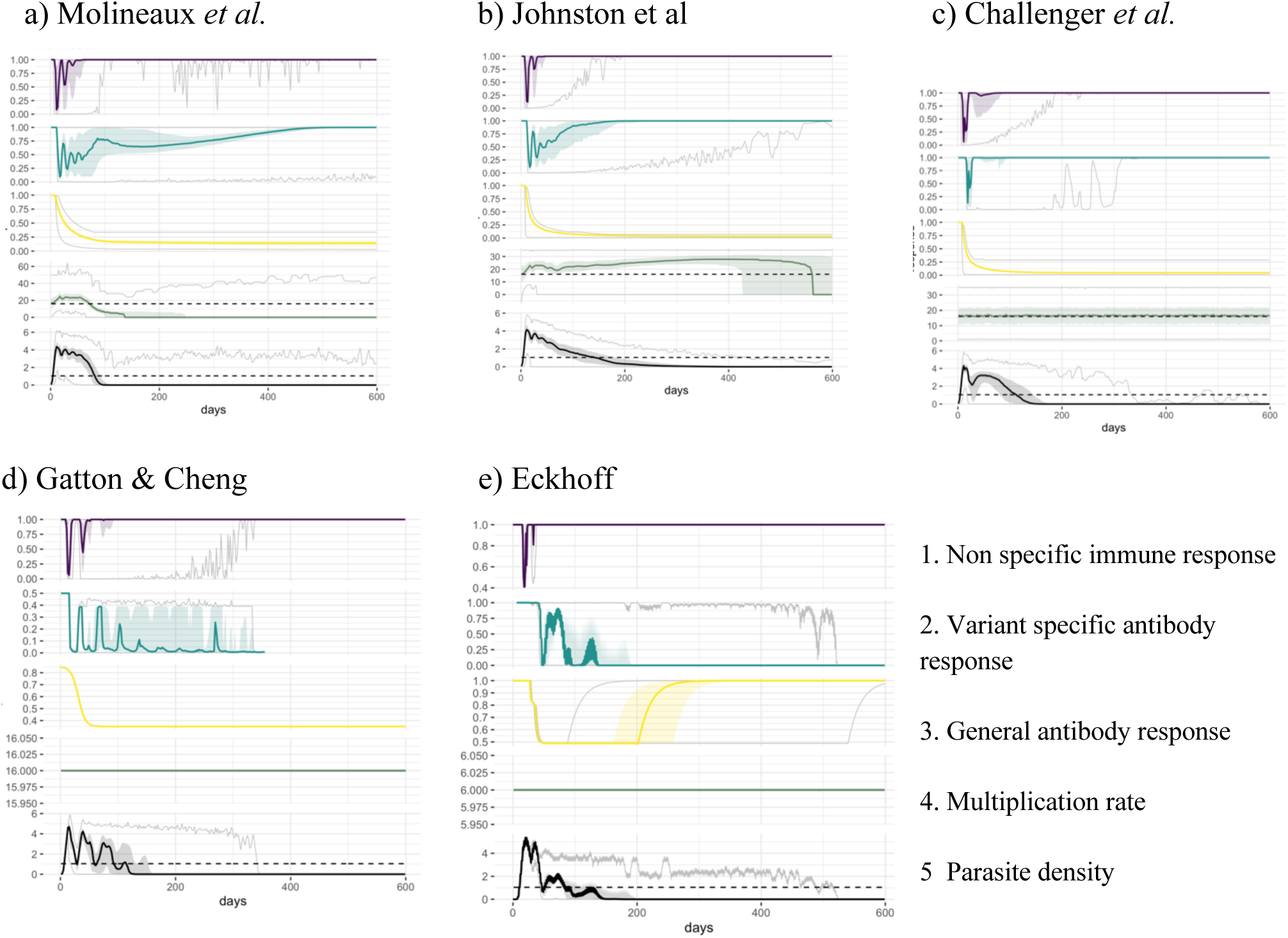
Immune responses, multiplication rates and asexual parasite densities in different models. Each panel shows, from top to bottom, 1. the effect of the innate immune response, 2. the effect of the variant specific immune response (see methods for calculations), 3. the effect of the general adaptive immune response, 4. the net effect of the product of the three immune responses, 5. the average (across variants, see methods) multiplication rate, 6. the log10 asexual density and 7. the product of the average multiplication rate with the net effect of the immune response. X-axis represents days of infection, solid line the median, shaded area the interquartile range (Q25-Q75) and the gray lines the minimum and maximum resulting from the 1750 simulations. The horizontal dashed green line represents an average multiplication rate of 16, the horizontal dashed black line indicate the threshold of positive density (10PBRC/μl). Results are shown for **a)** Molineaux et al., **b)** Johnston et al., and **c)** Challenger et al., **d)** Gatton & Cheng, and **e)** Eckhoff

**Table S2:**
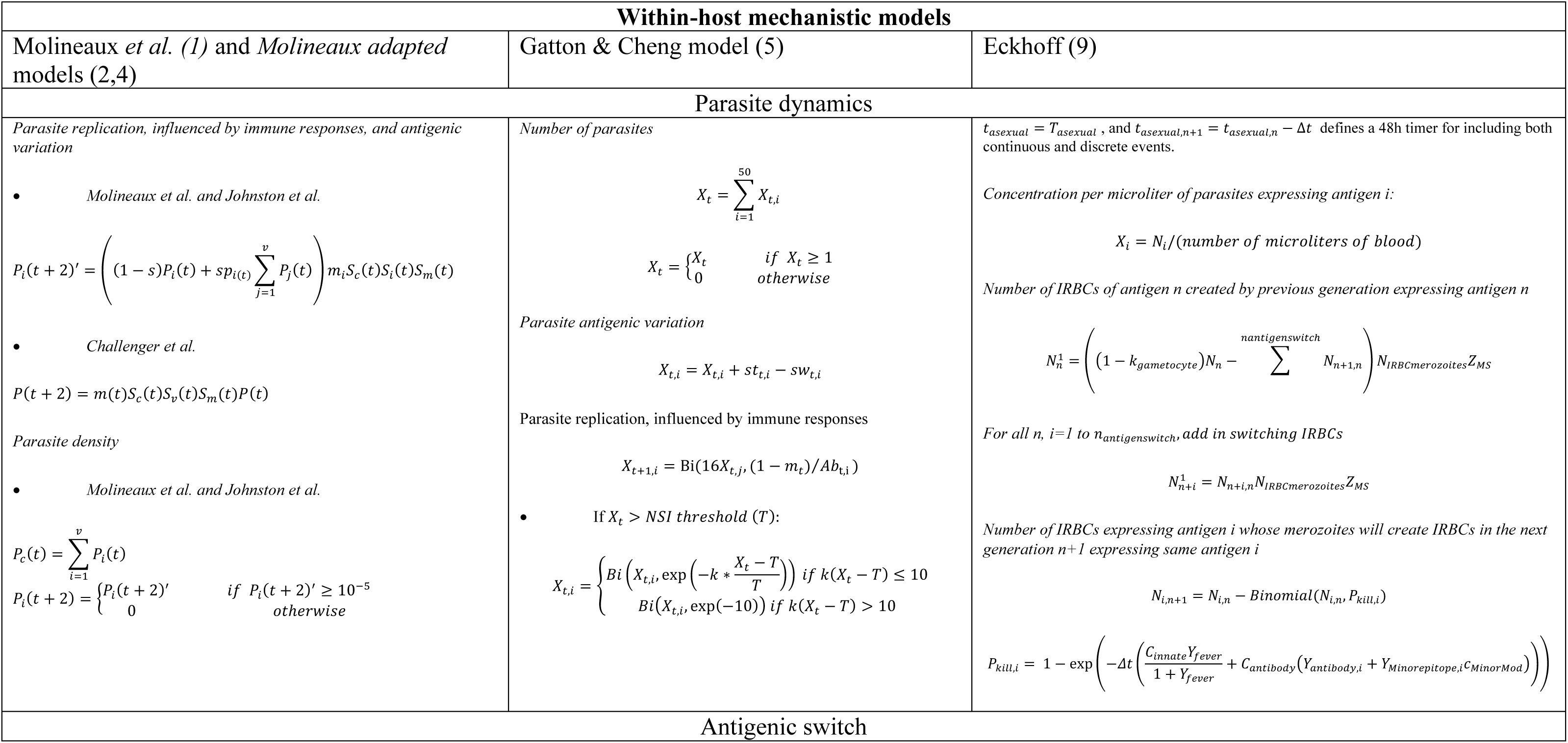
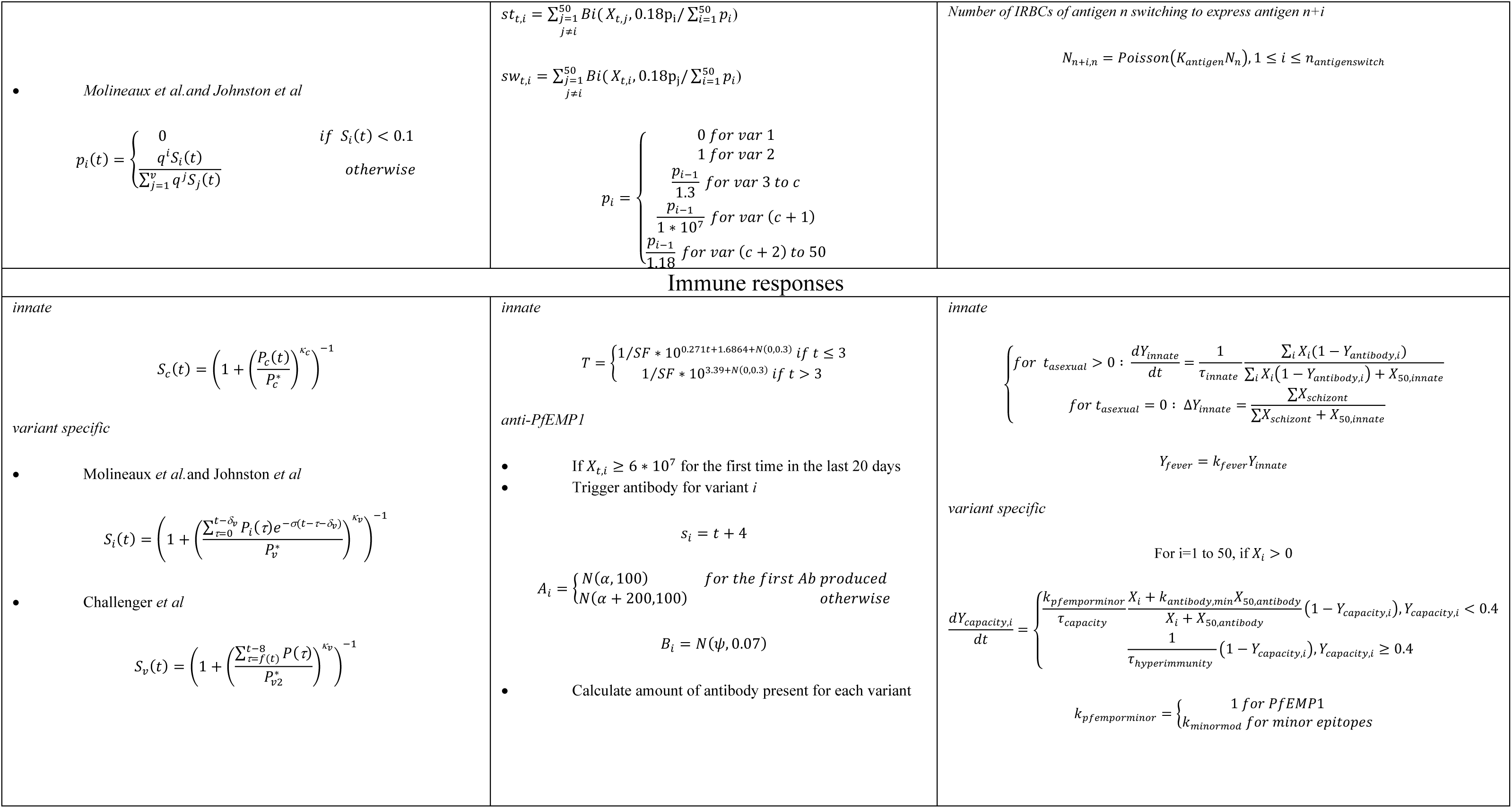
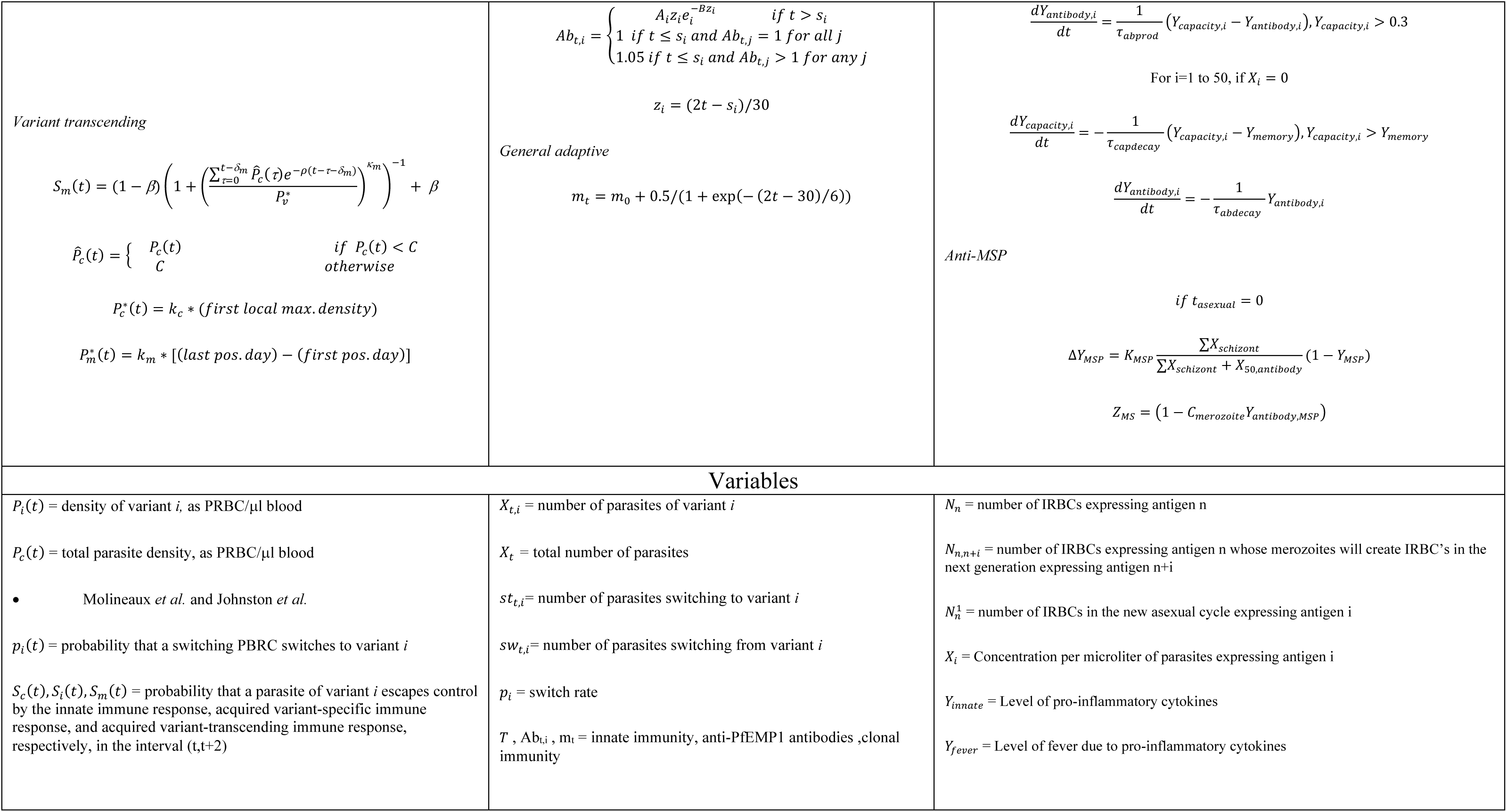
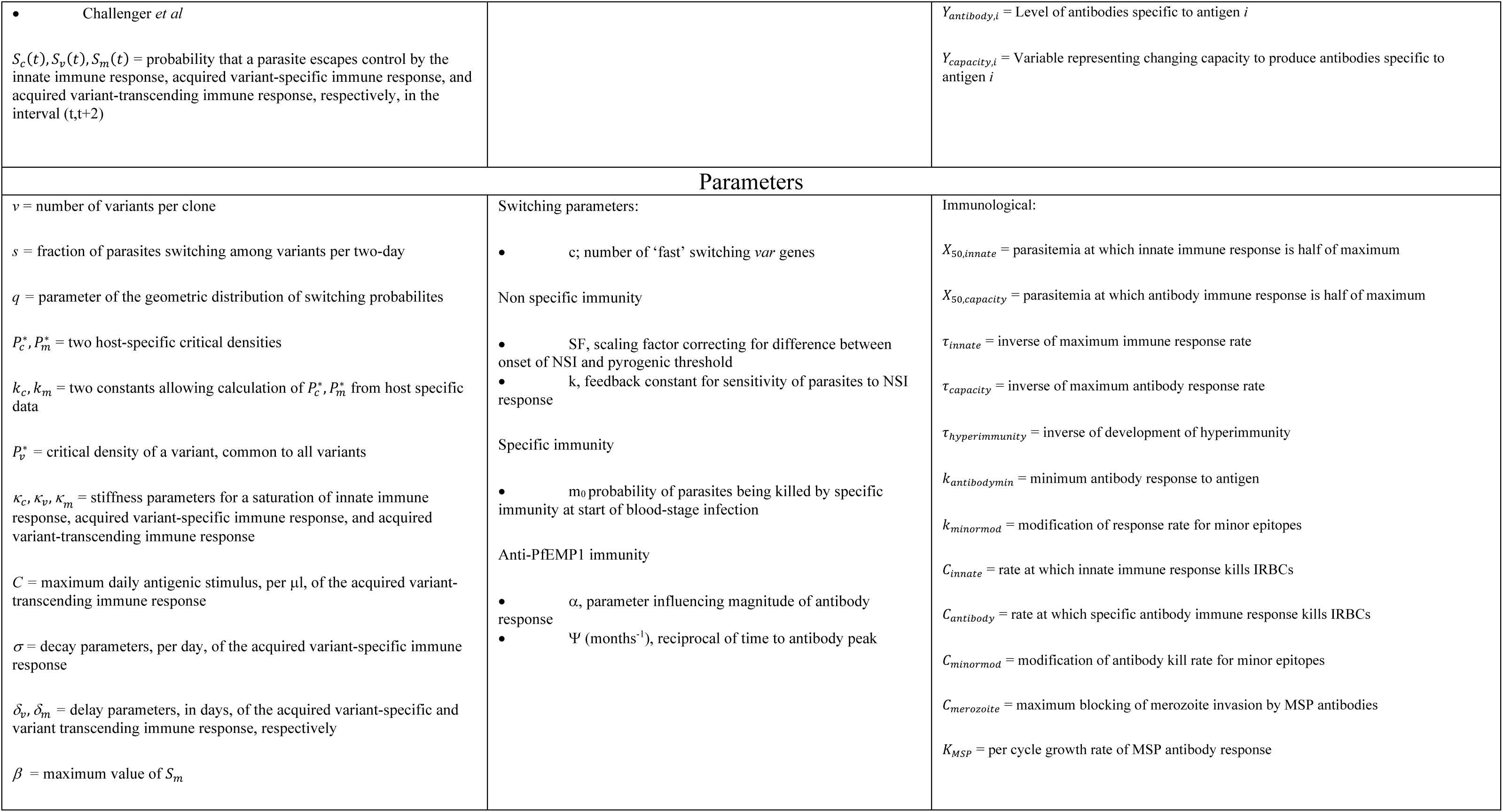
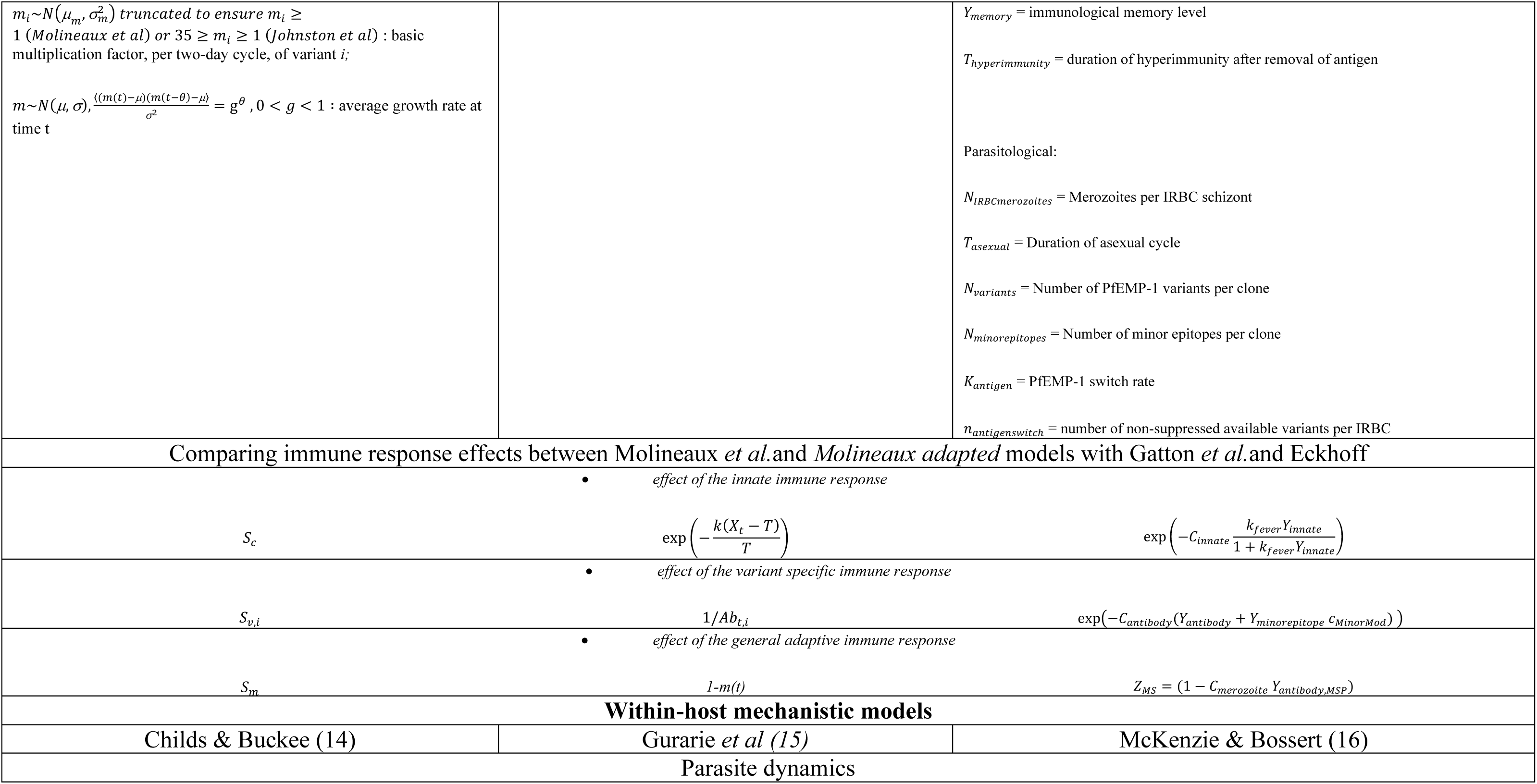
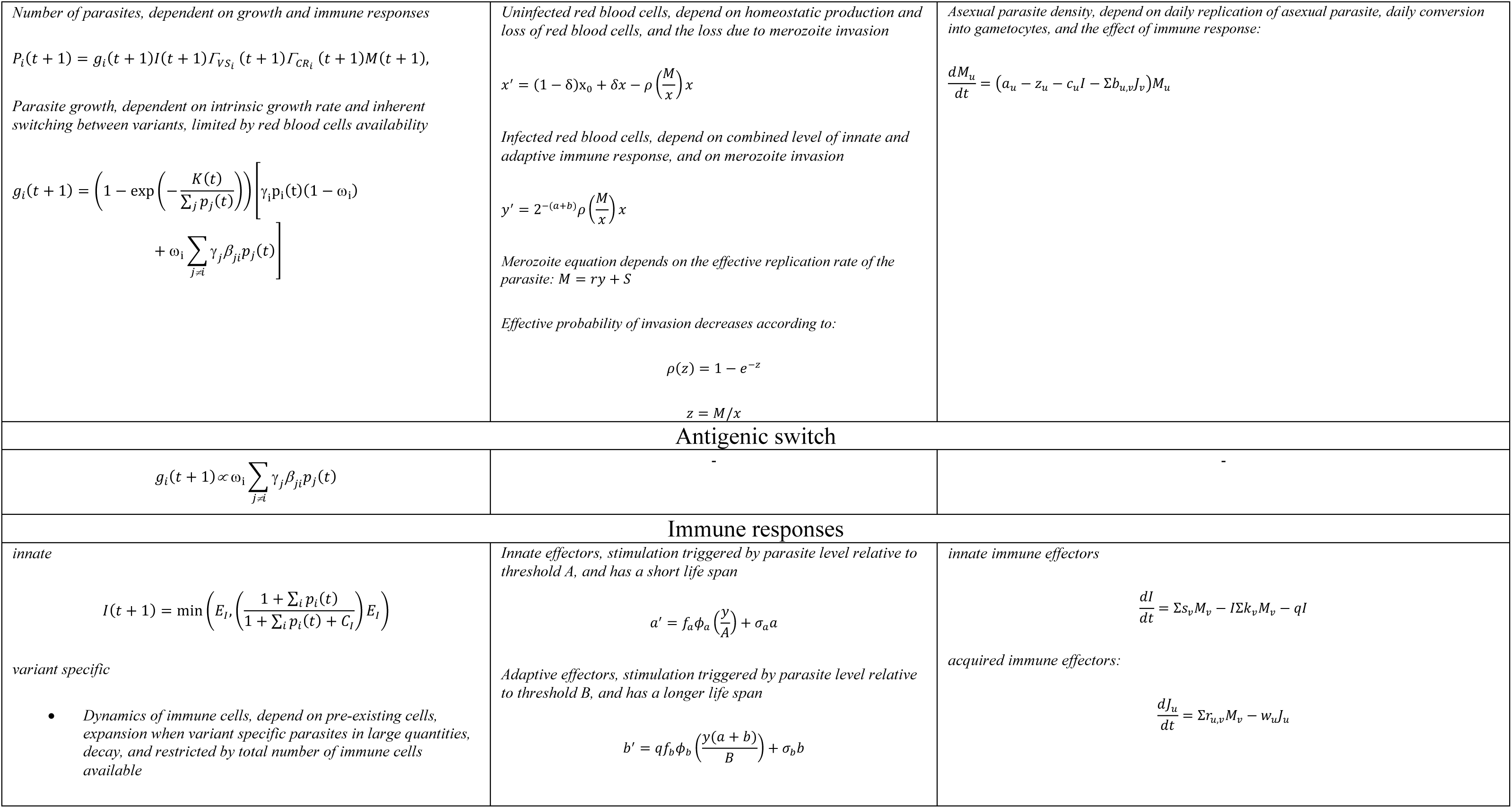
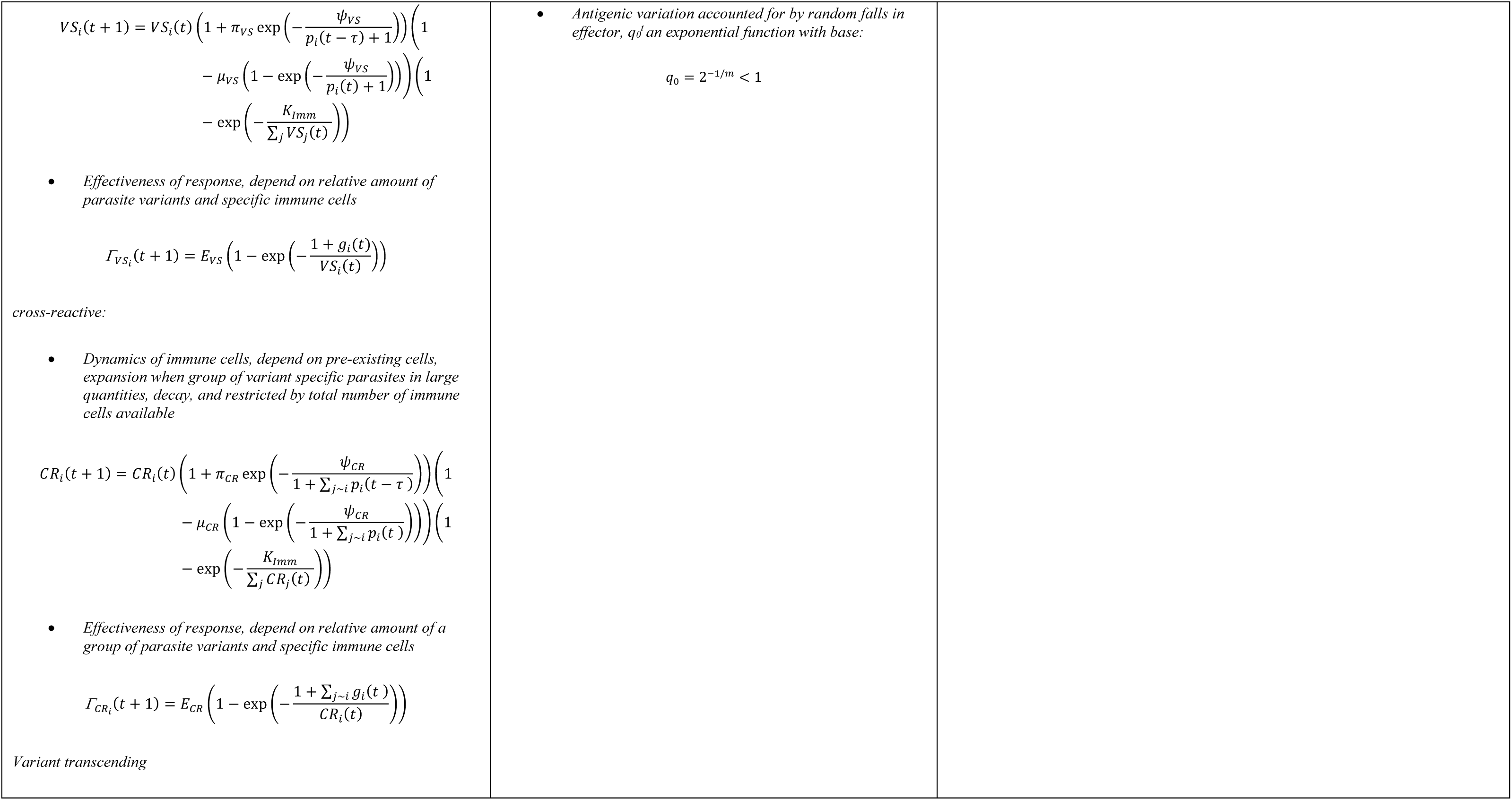
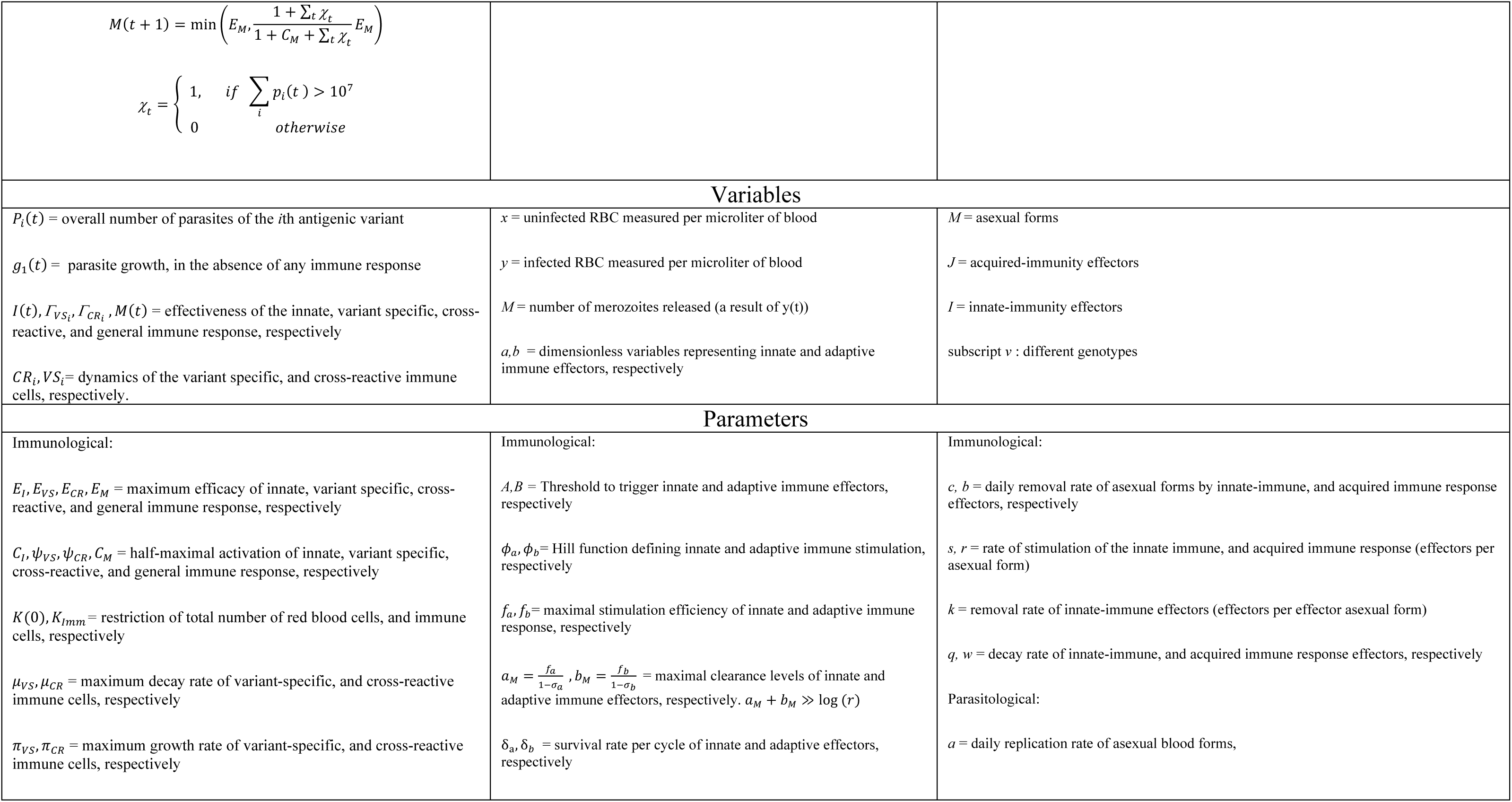
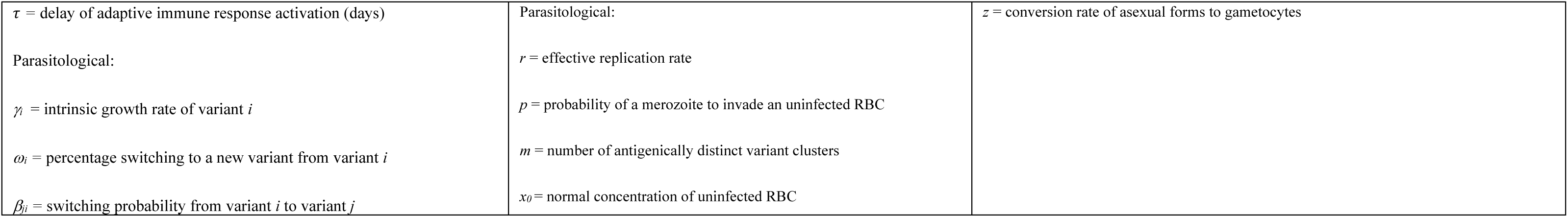
Model equations.

**Table S2:**
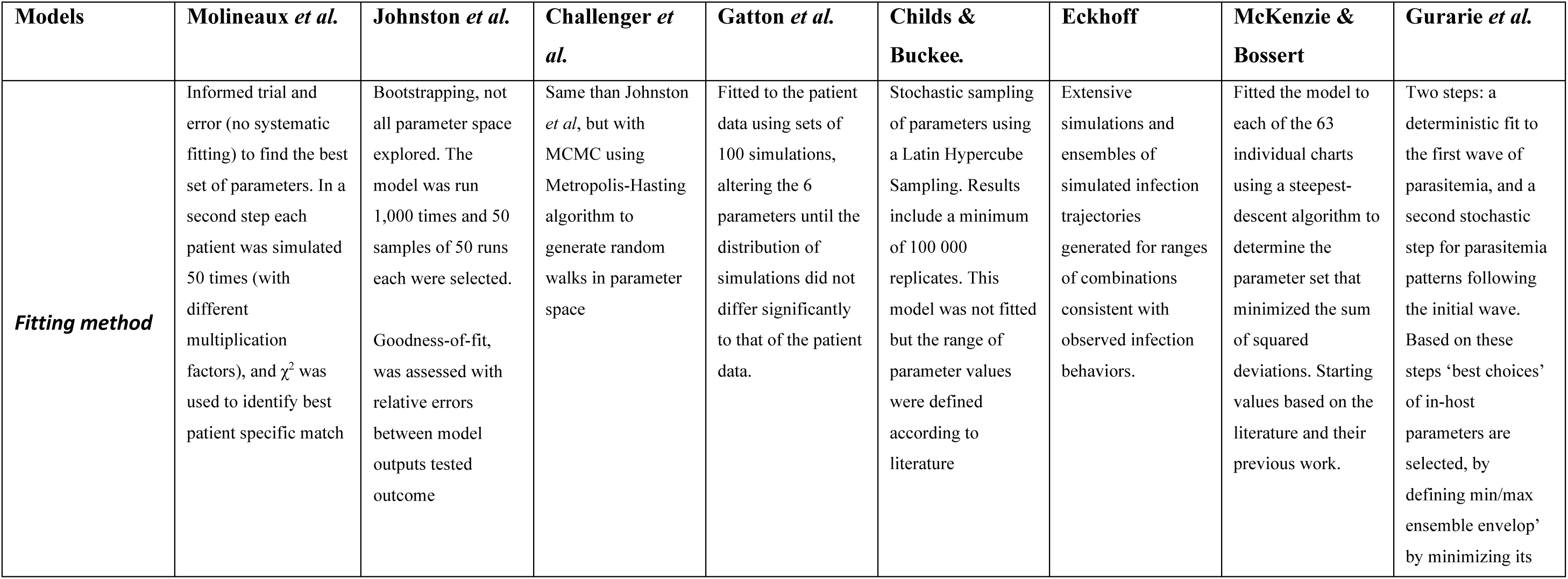
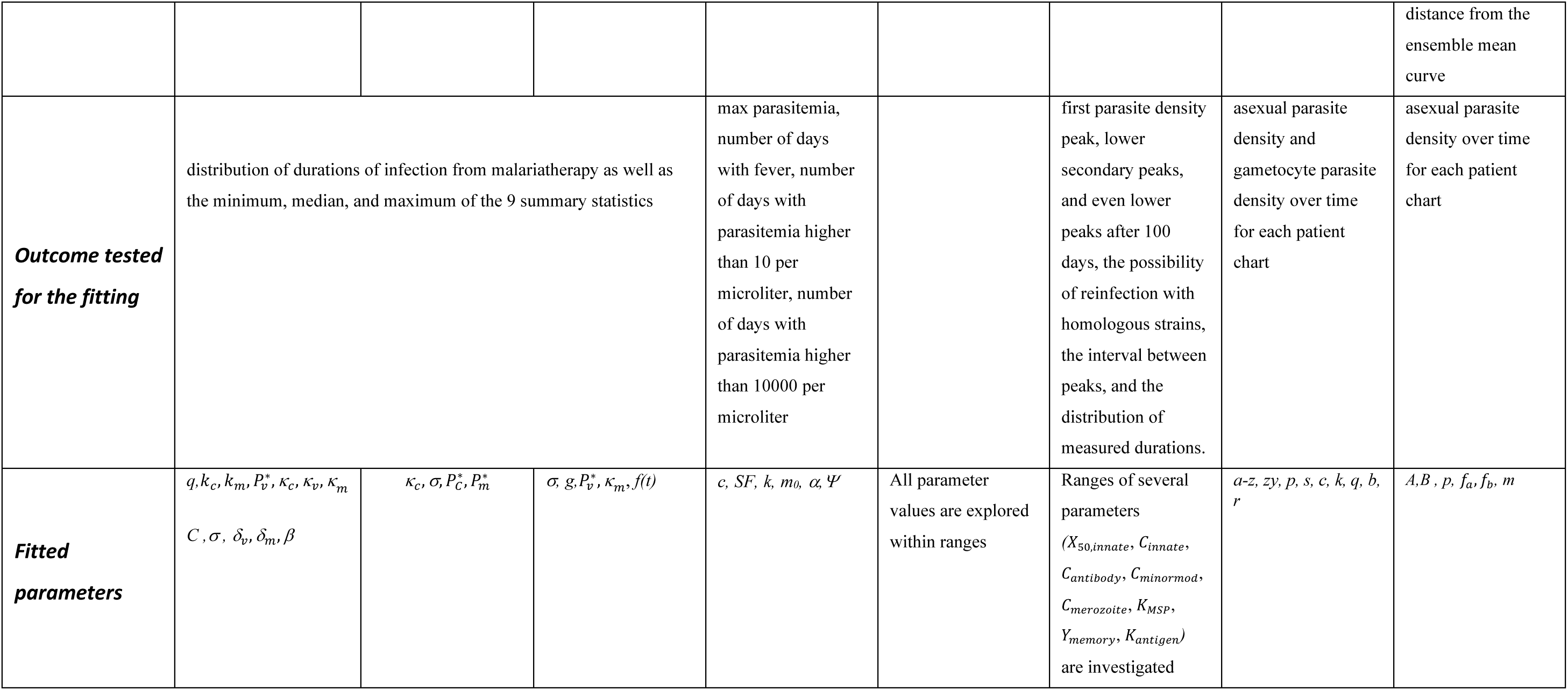
**Model fitting,** as described in the initial publications. Fitting methods and data might have been modified subsequently.

